# Role of Posterior Medial Thalamus in the Modulation of Striatal Circuitry and Choice Behavior

**DOI:** 10.1101/2024.03.21.586152

**Authors:** Alex J. Yonk, Ivan Linares-García, Logan Pasternak, Sofia E. Juliani, Mark A. Gradwell, Arlene J. George, David J. Margolis

**Affiliations:** Department of Cell Biology and Neuroscience, Rutgers, The State University of New Jersey, 604 Allison Road, Piscataway, NJ, 08854, USA

**Keywords:** Posteromedial Thalamus, Thalamostriatal Signaling, Synaptic Physiology, Sensorimotor Integration, Photometry, Optogenetics, PV Interneurons, Striatum, Spiny Projection Neurons, Behavioral Choice

## Abstract

The posterior medial (POm) thalamus is heavily interconnected with sensory and motor circuitry and is likely involved in behavioral modulation and sensorimotor integration. POm provides axonal projections to the dorsal striatum, a hotspot of sensorimotor processing, yet the role of POm-striatal projections has remained undetermined. Using optogenetics with slice electrophysiology, we found that POm provides robust synaptic input to direct and indirect pathway striatal spiny projection neurons (D1- and D2-SPNs, respectively) and parvalbumin-expressing fast spiking interneurons (PVs). During the performance of a whisker-based tactile discrimination task, POm-striatal projections displayed learning-related activation correlating with anticipatory, but not reward-related, pupil dilation. Inhibition of POm-striatal axons across learning caused slower reaction times and an increase in the number of training sessions for expert performance. Our data indicate that POm-striatal inputs provide a behaviorally relevant arousal-related signal, which may prime striatal circuitry for efficient integration of subsequent choice-related inputs.

## Introduction

The process of sensorimotor learning is underpinned by sensory perception and motor control.^1^ In the mouse whisker system, tactile sensations are acquired via active sensor (e.g., whisker) movement to obtain relevant environmental information and subsequent processing by the well-characterized primary somatosensory barrel cortex (S1) circuitry.^1,2^ This whisker-related information is transmitted from the periphery to S1 via two thalamic nuclei, ventral posterior medial (VPM) and posterior medial (POm), constituting the lemniscal and paralemniscal pathways, respectively.^3,4,5,6,7,8,9,10^ VPM reliably encodes fast-whisking components including self-motion and tactile information.^7,11,12,13,14^ Conversely, POm encodes phase-related whisking activity with relatively lower magnitude responses and higher response failure rates.^7,12,13,14,15,16,17,18^ Recent work has highlighted two behavior-related aspects of POm function: (1) activation during changes in behavioral state, especially related to sensory and nociceptive processing,^7,11,12,13,17,18,19,20,21^ and (2) driving learning-related plasticity at its cortical synapses.^22,23,24,25,26^

POm receives a plethora of inputs including glutamatergic (S1, primary motor cortex (M1), secondary somatosensory cortex, superior colliculus, and spinal trigeminal interpolaris), ^15,27,28,29,30,31,32,33,34^ GABAergic (ventral zona incerta, anterior pretectal nucleus, and thalamic reticular nucleus),^16,35,36,37,38,39,40,41,42^ and cholinergic (pedunculopontine and laterodorsal tegmental nuclei).^43,44,45,46^ Further, the stereotypical POm-cortical projection terminates in S1, specifically layers (L)1 and L5A,^4,47,48,49^ and has been studied in the context of driving cortical plasticity/perceptual learning.^22,23,24,25,26,50,51^ In addition to its cortical projection, POm axons pass through and collateralize with terminal synaptic boutons in both thalamic reticular nucleus and posterior dorsolateral striatum (pDLS) as they ascend towards cortex.^49,52,53,54,55,56,57,58^ Here, we focus on the POm-striatal projection as striatal circuitry modulation may have powerful effects on sensorimotor integration and behavior.^59^ However, POm’s influence over striatal microcircuitry and behavioral performance is unresolved.

The striatum is the predominant input nucleus of the basal ganglia and is predominantly composed of GABAergic spiny projection neurons (SPNs) expressing either D1 or D2 dopamine receptors,^60,61,62^ but it also contains a rich diversity of interneurons, such as parvalbumin-expressing (PV) fast-spiking interneurons that exert robust modulatory control over SPN output.^63,64,65,66,67^ Within this microcircuitry, the dorsal striatum integrates widespread convergent cortical and thalamic inputs that constitute part of the force driving normal striatal functioning.^68,69,70,71,72,73,74,75,76^ Notably, functionally-related cortical (S1 and M1) and thalamic (POm) inputs converge within shared striatal subregions.^73,74,77^ For example, M1 and S1 are heavily interconnected via reciprocal L2/3 and L5 connections,^78,79,80^ and their projections overlap within dorsal striatum and even onto the same neuron.^73,81,82,83,84,85,86^ While some studies treat striatal inputs as a uniform entity,^87,88^ they have been shown to differ anatomically,^73,74,76^ functionally,^89,90^ and behaviorally.^89,91,92^ Thus, the specific cortical and thalamic origin of striatal inputs likely has significance for understanding how the striatal circuitry integrates sensorimotor information to modulate behavior.

The most prominent thalamostriatal modulation occurs via parafascicular (Pf) thalamus.^90,93,94,95,96,97,98,99,100,101^ Pf is implicated in regulating action flexibility^102,103^ and contributing to the initiation and execution of learned sequences of movements^101,104^ through its robust innervation of striatal cholinergic interneurons.^95,87,88,90^ Conversely, despite direct comparisons to Pf,^56^ POm’s functional innervation pattern and subsequent influence over the striatal microcircuitry and choice behavior is undetermined.^59^ Here, we used *ex vivo* whole-cell recordings of identified D1-SPNs, D2-SPNs, and PV interneurons to assess the functional connectivity of POm-striatal projections, and *in vivo* fiber photometry and photoinactivation to identify the contribution of POm-striatal axonal activity on sensory-guided behavioral performance and learning.

## Results

### POm Equally Innervates Striatal Cell Types With Faster Latency In PV Interneurons

We stereotaxically injected pAAV-ChR2-EYFP unilaterally in POm, permitting channelrhodopsin (ChR2) expression in its thalamostriatal terminals to investigate the relative synaptic strength of POm inputs onto three identified striatal neurons (D1-SPNs, D2-SPNs, and PV interneurons; **Figure 1A-B**). The injection site was confirmed by verifying the stereotypical POm-cortical projection pattern (S1 L1 and L5a; **Figure 1C**).^4,25,26,47,49^ In acute *ex vivo* brain slices, neurons were targeted for patch clamp recordings within pDLS (AP from bregma = -0.34 to -1.22) corresponding with the POm-striatal axonal projection field (**Figure S1**).^56,105^ Striatal neurons were identified and targeted by crossing their respective Cre-recombinase mouse lines with tdTomato-expressing reporter mice and validating their intrinsic electrophysiological properties in response to hyperpolarizing and depolarizing current steps (**Figure S1A-D, G-L; see Methods**).^64,67,106^ Whole-cell current-clamp recordings were performed without inhibitory synaptic blockers to resemble natural physiological responses to optogenetic activation of POm inputs.^89^ After breaking in, patched cells were subjected to a standard set of protocols: (1) hyperpolarizing and depolarizing current steps to define intrinsic firing properties and optogenetic; and (2) single pulse (SP), (3) paired-pulse ratio (PPR), and (4) train stimulation to measure synaptic responses.

**Figure 1.**
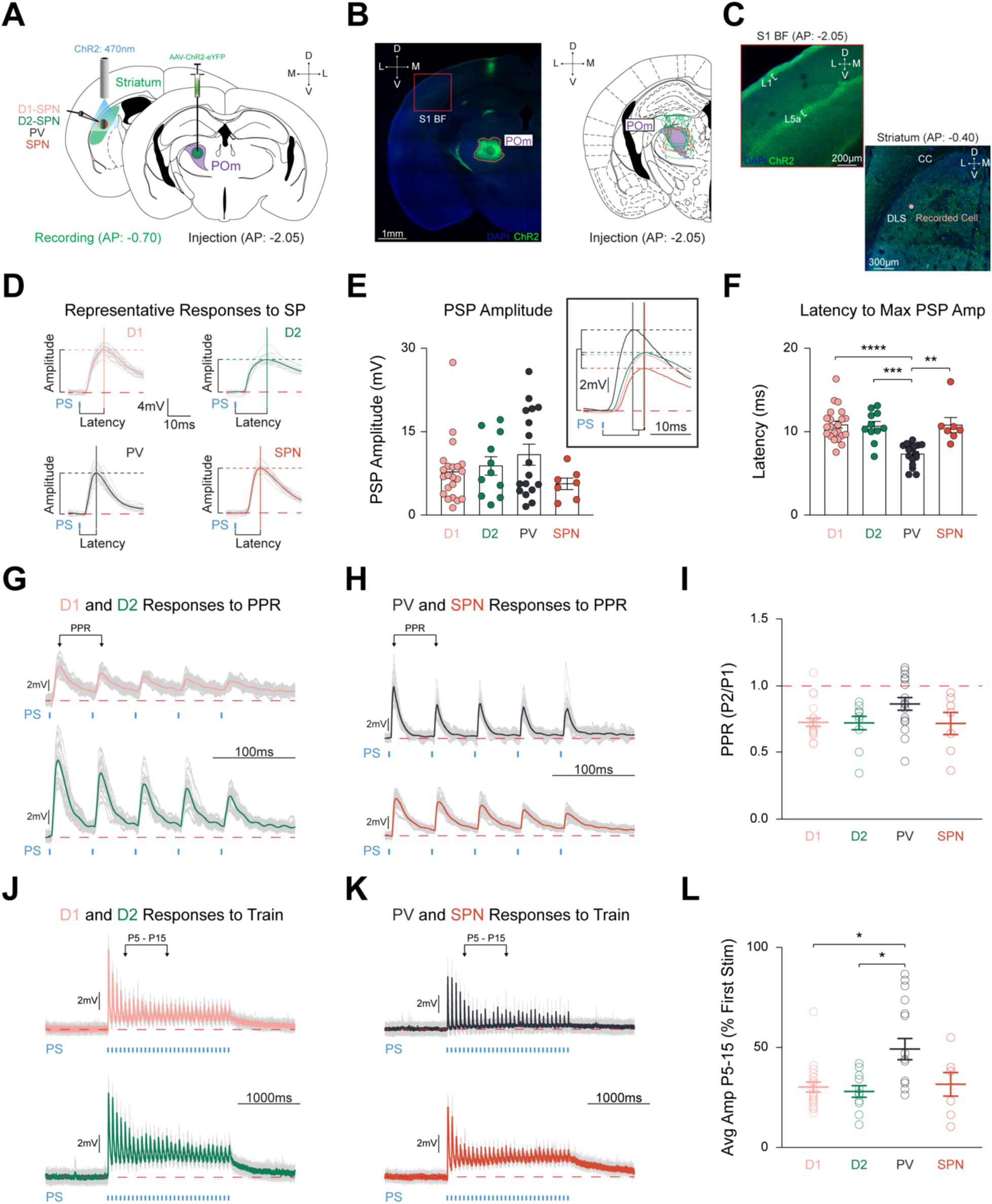
POm Equally Innervates Striatal Cell Types with Faster Latency In PV Interneurons. **(A)** Schematic detailing pAAV-ChR2-EYFP injection unilaterally into POm (*Right*), and optogenetic stimulation of POm-striatal afferents whilst recording from identified and unidentified neurons via *ex vivo* slice of posterior DLS (AP range: -0.34 to -1.22 relative to Bregma; *Left*). See **Figure S1**. Illumination (2.5ms pulses of 470nm light, ∼0.6mW intensity) was delivered through the 40x objective. **(B)** Representative injection site (orange) in POm (*Left*), and viral spread of all electrophysiology injections within highlighted POm (purple; *Right*). S1BF = S1 Barrel Field. Scale = 1mm. **(C)** Red box inset from panel **(B)** highlighting stereotypical POm-cortical projection pattern to S1BF L1 and L5a.^25,26,49^ *Right*: POm-striatal axons within posterior DLS. CC = corpus callosum. Scale = 200µm. **(D)** Representative cell type-specific PSPs to SP stimulation. Colored lines = average PSP of 20 sweeps. Gray lines = 20 individual traces. Solid vertical and dashed horizontal lines = latency and amplitude, respectively. Red dashed line = 0mV. Blue tick = photostimulation (PS). Time scale = 10ms. Voltage scale = 4mV. **(E)** Amplitudes evoked by each cell type were similar (D1-SPNs = 20 cells from 6 mice, D2-SPNs = 11 cells from 5 mice, PVs = 17 cells from 7 mice, unidentified SPNs = 7 cells from 4 mice). Inset shows grand average PSPs. Time scale = 10ms. Voltage scale = 2mV. **(F)** Latency to maximum PSP amplitude is significantly quicker in PVs than all other cell types. **(G-H)** Representative responses of **(G)** D1-SPN (*Top*) and D2-SPN (*Bottom*), and **(H)** PV (*Top*) and putative SPN (*Bottom*) to PPR stimulation.^109^ PPR is defined as the ratio of PSP amplitude of pulse 2 over the ratio of PSP amplitude of pulse 1. PPR PS parameters = five 2.5ms pulses with 50ms interpulse intervals (20Hz). Time scale = 100ms. Voltage scale = 2mV. See **Figure S2I)** Stimulation of POm-striatal afferents evokes similar PPR responses. **(J-K)** Representative responses of **(J)** D1-SPN (*Top*) and D2-SPN (*Bottom*), and **(K)** PV (*Top*) and putative SPN (*Bottom*) to train stimulation.^111^ Colored lines = average of 5 individual gray traces. Train PS parameters = thirty 2.5ms pulses with 64.2ms interpulse intervals (15Hz). Time scale = 1000ms. Voltage scale = 2mV. **(L)** Relative PSP amplitude (average of pulses 5-15 compared to pulse 1) is significantly larger than both SPNs. Data are mean ± SEM. *p < 0.05, ** p < 0.01, *** p < 0.001, **** p < 0.0001.

Optogenetic activation of POm terminals readily elicited depolarizing postsynaptic potentials (PSP) in all targeted cell types (**Figure 1D**). Responses to SP stimulation resulted in relatively equal PSP amplitudes for D1-SPNs (7.05±0.75mV), D2-SPNs (8.79±1.67mV), PVs (10.83±1.91mV), and neighboring unlabeled cells that we termed putative SPNs, based on their intrinsic firing properties (5.54±1.04mV) (F_(4,51)_ = 2.455, p = 0.4835; **Figure 1E**). Three PV interneurons and one D1-SPN exhibited action potentials to SP stimulation and were excluded from further analysis. A small but significant correlation was observed between PSP amplitude and increasing distance from the injection site (r^2^ = 0.08493, p = 0.0293, n = 59 cells; **Figure S1E-F**). The latency to maximum PSP amplitude was significantly shorter in PVs (7.29±0.32ms, n = 17 cells from 7 mice) than D1-SPNs (11.05±0.45ms, n = 20 cells from 6 mice), D2-SPNs (10.68±0.56ms, n = 11 cells from 5 mice), and putative SPNs (10.81±0.92ms, n = 7 cells from 4 mice) (F_(4,51)_ = 29.78, p < 0.0001, PV vs. D1 p < 0.0001, PV vs. D2 p = 0.0003, PV vs. SPN, p = 0.0056; **Figure 1F**). In a subset of recordings, identified and unidentified cells within the same field of view on the same slice were patched sequentially to control for injection site variability. D1- and D2-SPNs did not differ in PSP amplitude (9.55±2.83mV in D1-SPNs vs. 8.48±2.15mV in D2-SPNs, p = 0.6406, n = 8 pairs, N = 5) or latency (10.13±0.66ms in D1-SPNs vs. 10.92±0.56ms in D2-SPNs, p = 0.3254, n = 8 pairs, N = 5; **Figure S1M-O**). In contrast, sequentially patched PV and SPNs did not significantly differ in maximum PSP amplitude (10.26±3.05mV in PVs vs. 4.76±0.97mV in SPNs, p = 0.2500, n = 9 pairs, N = 7), but PVs had faster latency (6.85±0.42ms in PVs vs. 10.75±0.87ms in SPNs, p = 0.0006, n = 9 pairs, N = 7; **Figure S1P-R**), validating the population results. Strikingly, we found relatively equal PSP amplitudes in all recorded cell types within pDLS, indicating that POm provides robust and unbiased synaptic input to all targeted striatal cells.

### Short-Term Synaptic Dynamics Are Similar Between Striatal Cell Types, But Synaptic Depression Is Milder in PV Interneurons

The strength of synaptic inputs varies dramatically based on activation frequency and in a cell-type-specific manner with robust synaptic contacts generally exhibiting synaptic depression.^89,90,107,108^ To fully characterize the relative synaptic strength of POm-striatal inputs in a cell-type-specific manner, we assessed short-term plasticity by applying a PPR protocol of five pulses (**Figures 1G-H, S2A-B)**.^109,110^ While all cell types exhibited robust synaptic depression overall, no PPR differences were observed (D1-SPNs: 0.73±0.03, n = 20, D2-SPNs: 0.72±0.05, n = 11, PVs: 0.86±0.05, n = 17, SPNs: 0.71 ± 0.08, n = 7) (F_(4,51)_ = 7.101, p = 0.0688; **Figure 1I**).

To further characterize short-term synaptic dynamics, we applied a train protocol of thirty pulses at a frequency characteristic of POm-striatal activity (**Figure 1J-K**).^18,111^ Similar to PPR stimulation, train stimulation elicited overall synaptic depression in all cell types, but PV interneurons exhibited a milder synaptic depression relative to both SPN types that occurred predominantly between pulses 5-15 (**Figure 1L, S2**). PV interneurons (0.49±0.05, n = 17) showed significant differences compared to D1-SPNs (0.30±0.03, n = 20) and D2-SPNs (0.28±0.03, n = 11), but not with SPNs (0.31±0.06) (F_(4,51)_ = 10.99, p = 0.0118, D1-SPN vs. PV p = 0.0184, D2-SPN vs. PV p = 0.0481, PV vs. SPN p = 0.4940; **Figure 1L**). Thus, POm-striatal projections provide SPNs and PV interneurons with unbiased and large amplitude synaptic inputs, characterized by milder synaptic depression in PVs, highlighting a potentially significant role in modulating striatal microcircuitry.^89,90^

### Mice Rapidly Learn to Discriminate Between Two Textures

While specific sensorimotor integrative and learning roles have been proposed and tested for several striatal inputs,^69,72,87,89,90,104,112,113,114^ the role of POm-striatal projections is still unknown. To monitor the activation of POm-striatal projections, we injected pAAV-axon-jGCaMP8s unilaterally into left POm and implanted a 400µm core cannula into left pDLS (**Figure 2A-B**). Violet light (405nm) and blue light (470nm) were constantly delivered to pDLS throughout the entire session to measure the isosbestic and POm-pDLS axonal calcium signals, respectively. Using a similar protocol from our previous publication,^89^ water-restricted wild-type mice were trained on a whisker-based discrimination (Go/NoGo) paradigm. Mice received water for licking correctly (Hit) to the Go texture (P100 sandpaper). They received a white noise tone and a 12s time-out period for licking incorrectly (False Alarm; FA) to the NoGo texture (P1200 sandpaper; **Figure 2C**). If mice did not lick to the Go or NoGo texture, the texture retreated to its starting point, and trials were considered Miss and Correct Rejection (CR), respectively (**Figure 2D**). Additionally, pupil dynamics are a known metric of arousal,^115,116^ correlate well with POm activity,^18,26^ and exhibit outcome-dependent differences during the Go/NoGo paradigm.^117^ Therefore, synchronized orofacial video was captured during behavioral performance, and deep-learning^118,119^ pupillometry^120^ was applied to assess pupil dynamics during task performance (**Figure S3B**) in which our results mirrored previously reported outcome-dependent differences.^117^

**Figure 2.**
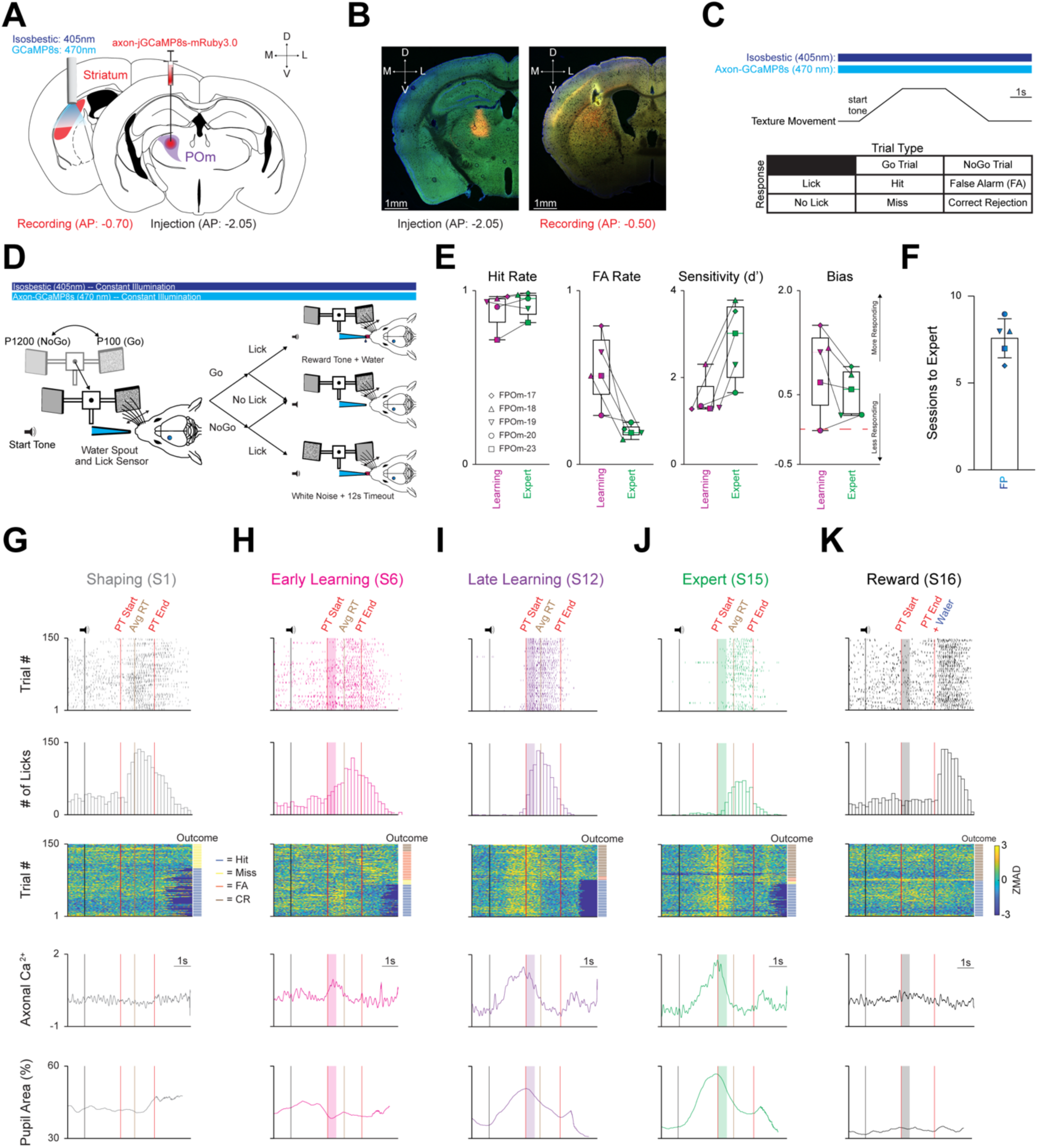
Mice Rapidly Learn to Discriminate Between Two Textures, and All Three Activity Parameters Markedly Increase Across Learning. **(A)** Schematic detailing pAAV-hSynapsin1-axon-jGCaMP8s-P2A-mRuby3 injection unilaterally into POm (*Right*), and a 400µm cannula implanted in the left posterior DLS (*Left*). **(B)** Representative injection site in POm (*Left*), and cannula placement in the posterior DLS along with ascending POm axons (*Right*). Scale = 1mm. **(C)** *Top*: Stimulating timing and texture movement representation during a trial. Note = both LEDs (isosbestic = 405nm; axon-jGCaMP8s = 470nm) were constantly on for every session. *Bottom*: Outcomes for each stimulus-response pair. **(D)** Schematic representing texture movement and potential outcomes during a single trial of the Go/NoGo whisker discrimination paradigm. **(E)** Changes in Hit Rate, FA Rate, Sensitivity (d’) and Bias of the FP cohort (n = 5 mice) as they transition from the Learning to the Expert phase. Note that mice are classified as Expert when they achieve a Hit Rate ≥ 0.80 and a FA Rate ≤ 0.30 for two consecutive sessions. Red line = 0. See **Figure S3**. **(F)** Average number of sessions required for expert discrimination of the FP cohort. **(G-K)** Three activity parameters (licking, axonal calcium, and pupil activity) from a representative **(G)** Shaping (session 1), **(H)** Early Learning (first two sessions after Shaping), **(I)** Late Learning (last two sessions before Expert), **(J)** Expert, and **(K)** Reward sessions from the same mouse (FPOm-18). *Top*: licking activity within a session (150 trials). Colored ticks = lick. Vertical black line = sound cue representing trial start as the texture moves towards the whisker field. Vertical red lines = start (texture arrival at endpoint in whisker field) and end (texture departure towards starting point) of the PT window (time where mice can respond by licking). Vertical brown line = average reaction time (RT; time of first lick that triggers an outcome) across all trials in each session. Colored boxes = 500ms grace period (licking does not trigger any outcomes). Note = no response line is present in the Reward session **(K)** as licking does not trigger any outcomes, and water was automatically delivered at PT end. *Top Middle*: Lick histogram. *Middle*: Heatmap sorted by trial outcome (to the *Right* of heatmap) highlighting axonal ZMAD calcium activity for each trial. Trial outcome is color coded (blue = Hit, yellow = Miss, orange = FA, brown = CR). *Bottom Middle*: Average axonal calcium activity of 150 trials for each session. *Bottom*: average pupil area (as a percentage) of 150 trials for each session. Data are mean ± SEM. Time scale = 1s.

Mice in the fiber photometry (FP) cohort (n = 5) underwent three training phases (Shaping, Learning, and Expert; **Figure S3A**) that were segmented into five discrete behavioral time points (Shaping, Early Learning, Late Learning, Expert, and Reward; **see Methods**). During the learning phase, Hit rate increased, and FA rate decreased significantly, leading to markedly increased sensitivity (d’) and slightly decreased bias (**Figure 2E, S3A**). Mice were considered Expert once they had reached ≥ 0.80 Hit Rate and ≤ 0.30 FA Rate for two consecutive sessions in lieu of a strict sensitivity (d’) threshold; we found this definition more intuitive because d’ is enhanced as Hit Rate and FA Rate approach their extremes (0 or 1) (**Figure S3A**). On average, it took this cohort 7.6±0.51 sessions to become Experts (**Figure 2F**). Thus, the FP cohort rapidly learned to discriminate between the two textures as validated by behavioral responding parameters.

### Calcium Activity Markedly Increases and Becomes Stereotyped Across Learning

To elucidate the contribution of POm-striatal projections during the Go/NoGo discrimination task, we measured POm axonal calcium activity along with licking and pupil activity at five discrete behavioral time points (Shaping, Early Learning, Late Learning, Expert, and Reward; **Figure 2G-K**). Photometry measurements revealed learning-related increases in POm axonal activity, starting before and peaking near texture presentation (**Figures 3A, S3A**) as measured by two parameters: calcium signal amplitude (ZMAD; median absolute deviation of the z-score) and area under the curve of the receiver-operator characteristic (auROC). First, the average maximum calcium amplitude significantly increased across learning (F_(1.818,7.271)_ = 39.24, p = 0.0001, n = 5; **Figure 3B**). Further, the average maximum amplitude at Shaping (0.27±0.04) was significantly smaller compared to Early Learning (1.04±0.12, p = 0.0190), Late Learning (1.58±0.18, p = 0.0061), and Expert (1.59±0.17, p = 0.0052). To test whether POm activity was reward-related, a single session was performed following the Expert phase during which the textures were removed, and water was automatically provided at the end of the presentation time (PT) window. We observed that calcium activity regressed to Shaping levels and was significantly smaller than Early (p = 0.0306), Late (p = 0.0126), and Expert (p = 0.0098).

**Figure 3.**
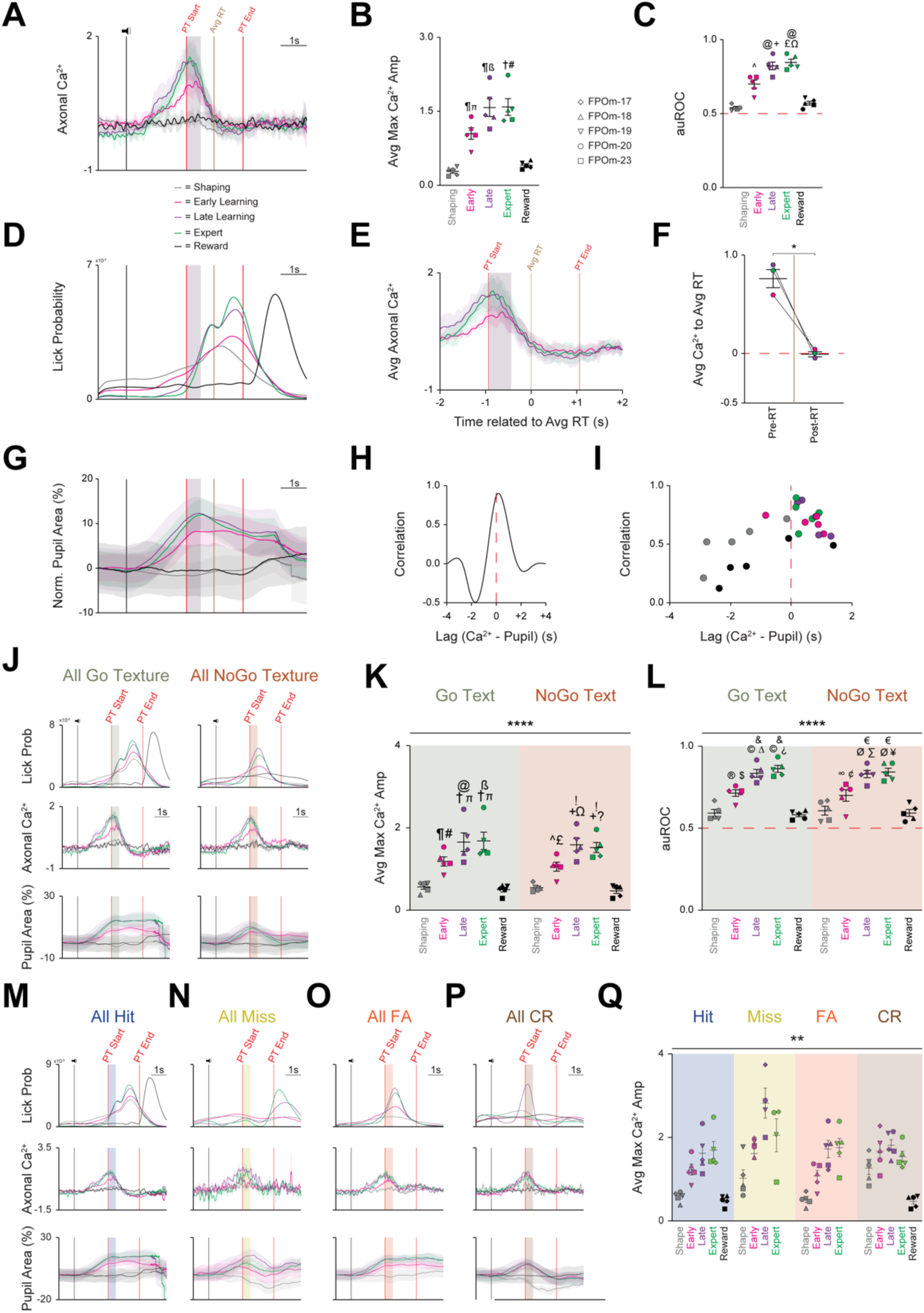
All Three Activity Parameters Exhibit Marked Increases Across Learning, But Only Axonal Calcium Activity Remains Unchanged, Irrespective of Trial Type or Outcome Segmentation. **(A)** Grand average axonal calcium activity at each behavioral time point. Data are mean ± SD. **(B)** Average of maximal axonal calcium amplitude markedly increases across learning before regressing to Shaping levels during the Reward session. ᴨ p < 0.01 Shaping vs. Early, ß p < 0.001 Shaping vs. Late, # p < 0.001 Shaping vs. Expert, ¶ p < 0.05 Early/Late vs. Reward, † p < 0.001 Expert vs. Reward. **(C)** Average area under the curve of the receiver-operator characteristic (auROC) also markedly increases across learning before regressing to Shaping levels during the Reward session. ^ p < 0.05, Shaping vs. Early; + p < 0.01, Shaping vs. Late; £ p < 0.001, Shaping vs. Expert; Ω p < 0.05, Early vs. Expert; @ p < 0.01, Late/Expert vs. Reward. **(D)** Grand average probability density function for licking-related activity at each time point. **(E)** Axonal calcium activity at Learning and Expert time points 2s pre and 2s post grand average RT. Data are mean ± SD. **(F)** Pre-RT axonal calcium activity is significantly larger than post-RT axonal calcium activity. **(G)** Grand average of normalized pupil area at each behavioral time point. **(H)** Representative cross-correlation of pupil area and axonal calcium activity**I)** Cross-correlation of pupil area and calcium activity plotted for each behavioral time point for each mouse. **(J)** Grand average of all licking (*Top*), calcium (*Middle*), and normalized pupil (*Bottom*) activity segmented by trial type: Go texture (*Left*) and NoGo texture (*Right*) presentation. **(K)** Average of maximal axonal calcium amplitude markedly increases across learning for both Go and NoGo texture presentation before regressing to Shaping levels during the Reward session. ¶ p = 0.0016 Go Shaping vs. Go Early, ᴨ p < 0.0001 Go Shaping vs. Go Late/Expert, @ p = 0.0265 Go Early vs. Go Late, ß p = 0.0154 Go Early vs. Go Expert, # p = 0.0005 Go Early vs. Go Reward, † p < 0.0001 Go Late/Expert vs. Go Reward. ^ p = 0.0128 NoGo Shape vs. NoGo Early, + p < 0.0001, NoGo Shape vs. NoGo Late/Expert, £ p = 0.0031 NoGo Early vs. NoGo Reward, Ω p = 0.0085 NoGo Early vs. NoGo Late,p = 0.0295 NoGo Early vs. NoGo Expert, ! p < 0.0001, NoGo Late/Expert vs. NoGo Reward. **(L)** Average auROC markedly increases across learning for both Go and NoGo texture presentation before regressing to Shaping levels during the Reward session. ® p = 0.0041 Go Shaping vs. Go Early, © p < 0.0001 Go Shaping vs. Go Late/Expert, Δ p = 0.0041 Go Early vs. Go Late, ¿ p = 0.0003 Go Early vs. Go Expert, $ p = 0.0014 Go Early vs. Go Reward, & p < 0.0001 Go Late/Expert vs. Go Reward. ∞ p = 0.0462 NoGo Shaping vs. NoGo Early, Ø p < 0.0001 NoGo Shaping vs. NoGo Late/Expert, ∑ p = 0.0017 NoGo Early vs. NoGo Late, ¥ p = 0.0005 NoGo Early vs. NoGo Expert, ¢ p = 0.0142 NoGo Early vs. NoGo Reward, € p < 0.0001 NoGo Late/Expert vs. NoGo Reward. **(M-P)** Grand average of licking (*Top*), calcium (*Middle*), and normalized pupil (*Bottom*) activity segmented by trial outcomes: **(M)** Hit, **(N)** Miss, **(O)** FA, and **(P)** CR. **(Q)** Average of maximal axonal calcium amplitude of each mouse markedly increases across learning for all trials outcomes before regressing to Shaping levels during the Reward session. Data are mean ± SEM unless noted otherwise. * p < 0.05, ** p < 0.01, **** p < 0.0001. See also **Figure S4**.

Second, to quantify POm axonal activity within the striatum relative to learning, we employed signal detection theory, utilizing auROC values to compare the basal activity across the five behavioral time points as learning occurs.^121,122^ As with the maximum calcium amplitude, auROC values significantly increased across learning (F_(2.162,8.650)_ = 51.17, p < 0.0001, n = 5; **Figure 3C**). Notably, auROC values at Shaping (0.54±0.01) were significantly lower than Early Learning (0.70±0.03, p = 0.0116), Late Learning (0.83±0.03, p = 0.0017), and Expert (0.85±0.02, p = 0.0007), but not Reward (0.57±0.01, p = 0.4111). Further, significant auROC differences were also noted between the Early Learning vs. Expert (p = 0.0383), Late vs. Reward (p = 0.0084), and Expert vs. Reward (p = 0.0042). Finally, calcium-related events increased longitudinally and became stereotyped within a 4s target window (centered around texture presentation) compared to a non-task-related 1s control window (during the 1s pre-task interval prior to the sound cue and texture movement; **Figure S4A-D**). Thus, POm-striatal projections represent a learning-related signal that increases prior to and becomes stereotyped to texture presentation.

### Licking and Pupil Dilation Also Markedly Increase Across Learning, But Only Pupil Is Correlated with Calcium Activity

Both licking and pupil dilation also exhibited marked increases across learning (**Figures 2G-K, S3A-B**). Licks occurring before texture presentation decreased dramatically as licking became stereotyped within the PT window (**Figure 3D**). We assessed whether licking and POm activity were correlated as both increase across learning (**Figure 3A, D**) by plotting the average calcium activity from the Early, Late, and Expert time points with the grand average reaction time (RT) set to 0 (**Figure 3E**). Comparison of average calcium activity pre- and post-RT revealed an overall significant reduction (0.79±0.09 in pre-RT vs. 0.00±0.03 in post-RT, p = 0.0221; **Figure 3F**), indicating no correlation between POm and licking activity, validating previous results.^26^

Pupil dilation started immediately following the cue sound and peaked near texture presentation before slowly decreasing throughout the rest of the trial (**Figure 3G**). Similar to previous studies,^18,26^ we found that pupil and POm activity were tightly correlated but decoupled during the PT window with POm activity immediately regressing to baseline, while pupil activity remained elevated (**Figure 3A, G**). Pupil dilation lagged by ∼250ms on average (**Figure 3H**), and the correlation became more consistent across learning and was restricted to before and at texture presentation (**Figure 3I**). Thus, despite all three activity parameters increasing across learning, POm activity is not related to motor activity and is only correlated with pre-PT pupil dilation.

### Increased POm Axonal Calcium Activity Is Independent of Trial Type or Outcome

This whole-session analysis did not account for sensory (texture) or outcome differences. To assess whether POm axonal activity is sensory-related (texture-specific), sessions were segmented by the presented texture (Go or NoGo; **Figure S4E-F**) and by trial outcome (below). Notably, licking and pupil dynamics have discrete activation patterns based on the presented texture.^117^ Licking activity in the Go condition occurs squarely within the PT window and is indistinguishable from whole-session licking activity (**Figure 3D**), while licking peaks near the end of the grace period in the NoGo condition (**Figure 3J**). Similarly, pupil activity starts to increase at the same point in both conditions but deviates at the end of the grace period. Conversely, POm axonal activity remained strikingly consistent between the Go and NoGo conditions (**Figure 3J**). This observation was further validated by comparing the average maximum calcium amplitude of each behavioral time point between the presented texture conditions. The main effect of behavioral time points was significant (F _(4,32)_ = 57.16, p < 0.0001), but the main effect of presented texture was not (F _(1,8)_ = 0.3797, p = 0.5549; **Figure 3K**). Post-hoc comparisons found that the maximum calcium amplitude during Shaping (Go: 0.55±0.06; NoGo: 0.53±0.04) was significantly reduced compared to Early Learning (Go: 1.18±0.11, p = 0.0016; NoGo: 1.05±0.11, p = 0.0128), Late Learning (Go: 1.65±0.23, p < 0.0001; NoGo: 1.59±0.16, p < 0.0001), and Expert sessions (Go: 1.68±0.21, p < 0.0001; NoGo: 1.52±0.12, p < 0.0001), but not the Reward session. The same overall effects were noted for the auROC analysis (main effect of behavioral time point: F _(4,32)_ = 66.43, p < 0.0001; main effect of presented texture: F _(1,8)_ = 0.01694, p = 0.8997; **Figure 3L**). As with the maximum amplitude, post-hoc comparison highlighted that the auROC value during Shaping (Go: 0.59±0.02; NoGo: 0.61±0.03) was significantly smaller than Early Learning (Go: 0.72±0.02, p = 0.0035; NoGo: 0.70±0.04, p = 0.0355), Late Learning (Go: 0.84±0.02, p < 0.0001; NoGo: 0.83±0.02, p < 0.0001), and Expert sessions (Go: 0.87±0.02, p < 0.0001; NoGo: 0.85±0.02, p < 0.0001), but not the Reward session. Thus, despite divergent licking and pupil activity as a function of the presented texture, axonal calcium activity remained unchanged, indicating POm-striatal projections do not encode vibrissae-specific sensory information.

We next tested whether POm calcium activity correlated with trial outcome (e.g., Hit, Miss, False Alarm, Correct Rejection; **Figure 2C-D**). Upon segmentation by trial outcome (**Figure S4G-J**), licking and pupil dynamics exhibited discrete outcome-dependent activity patterns,^117^ but calcium activity again remained remarkably consistent (**Figure 3M-P**). These results were further validated by comparing the average maximum calcium amplitude of each behavioral time point between the outcome conditions: the main effects of behavioral time point (F _(1.273,5.090)_ = 22.09, p = 0.0043) and trial outcome (F_(1,153, 4.613)_ = 10.80, p = 0.0231; **Figure 3Q**) were significant. However, the only significant post-hoc comparisons were between Hit-Shaping and CR-Shaping (p = 0.0305) and FA-Shaping and CR-Shaping (p = 0.0114). Overall, POm-striatal projections do not appear to encode texture- or outcome-specific information, suggesting that POm may represent a behaviorally relevant arousal-related role.

### Optogenetic Suppression of POm-Striatal Axons During Behavior Slows Learning Rate

Our findings indicate that POm-striatal inputs may represent an arousal signal. To investigate this possibility, wild-type mice were divided into two cohorts: a No Stim and a photoinactivation JAWS cohort. For both cohorts, a 200µm fiber optic cannula was unilaterally implanted over left pDLS. However, only the JAWS cohort received an injection of the inhibitory actuator JAWS (**Figure 4A-B**)^123,124,125^ into ipsilateral POm, verified by the stereotypical POm-cortical projection pattern (**Figure 4B**).^4,47,49^ Both cohorts were trained on the Go/NoGo task (**Figures S3, 5**). However, once the Learning phase started, POm-striatal axons of the JAWS cohort were photoinactivated for ∼50% of trials per session until reaching the Expert phase. Photoinactivation occurred via an amber (617nm) LED (∼7mW) for 2s of constant illumination centered around the PT window start (**Figure 4C-D**) where the maximum calcium signals were detected (**Figure 3A**). Both the No Stim and JAWS cohorts achieved Expert status (**Figure S5A-B**) by increasing Hit Rate, decreasing FA Rate, and with licking becoming stereotyped across learning (**Figures 4E-H, S5A-B**). However, photoinactivation of POm-striatal axons during the Learning phase resulted in the JAWS cohort requiring significantly more sessions (12.50±0.50 sessions, n = 4 mice) to attain Expert status compared to the other two cohorts with different experimental conditions (FP with constant illumination across the entire session: 7.60±0.51 sessions, n = 5 mice; No Stim with no illumination: 7.40±0.24 sessions, n = 5 mice) (F_(3,11)_ = 8.542, p = 0.0041, JAWS vs. FP p = 0.0470; JAWS vs. No Stim p = 0.0202; FP vs. No Stim p > 0.9999; **Figure 4I-J**). We assessed whether photoinactivation modified behavioral responding parameters between the Learning and Expert phases (**Figure 3K**). The only significant effect was a decrease in overall responding (Bias) during the Learning phase (Off: 0.56±0.32, On: 0.48±0.32, p = 0.0351). Due to this, we assessed if inhibition resulted in slower RTs and found that the average RT was significantly slower in the On condition (Off: 0.87±0.07, On: 0.93±0.07, p = 0.0124). Thus, the suppression of this behaviorally relevant arousal signal resulted in more learning sessions due to delayed RTs.

**Figure 4.**
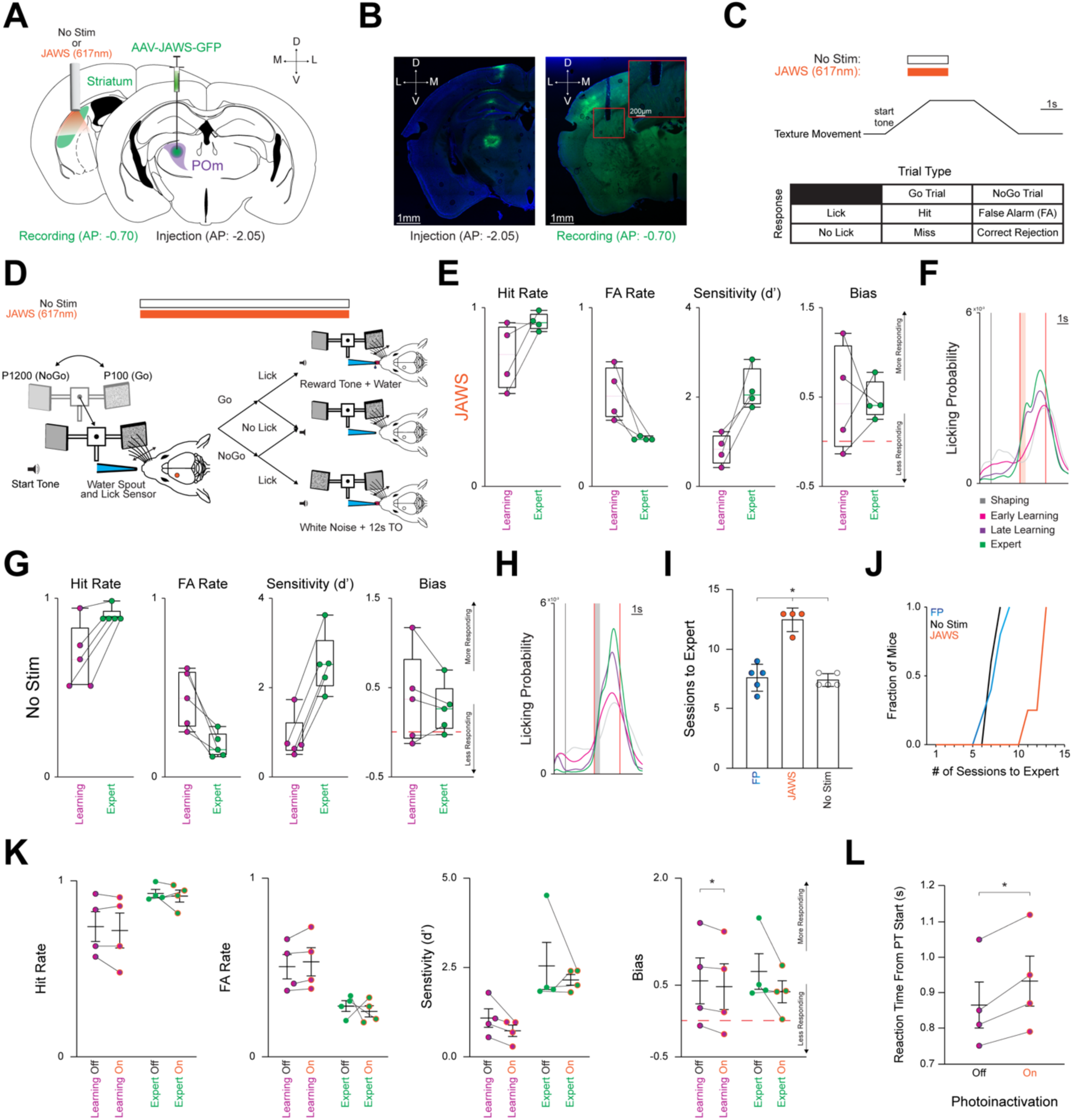
Photoinactivation Increases Number of Sessions Required For Expert Discrimination. **(A)** Schematic detailing pAAV-hSyn-JAWS-KGC-GFP-ER2 (JAWS) injection unilaterally into POm (*Right*) and a 200µm cannula implanted in the left posterior DLS (*Left*). For the No Stim cohort, only the cannula was implanted in the left posterior DLS. Activation of the inhibitory JAWS opsin was performed constantly on for 1s before and after texture arrival in the whisker field. JAWS activation probability per trial = 0.50. **(B)** Representative injection site in POm (*Left*), and the cannula placement in the posterior DLS along with ascending POm axons (*Right*). Scale = 1mm. Red inset shows ascending POm axons underneath the optic cannula. Inset scale = 200µm. **(C)** *Top:* Stimulation timing (constant illumination for 2s, centered around texture arriving at its endpoint) and texture movement representation during a trial. Note that no light is presented for the No Stim cohort as no stimulation occurred. *Bottom:* Outcomes for each stimulus-response pair. **(D)** Schematic representing texture movement and potential outcomes during a single trial. **(E)** Changes in Hit Rate, FA Rate, Sensitivity (d’), and Bias of all JAWS cohort mice (n = 4) as they transition from the Learning to the Expert phase in box-and-whisker plots. Note that mice are classified as Expert when they achieve a Hit Rate ≥ 0.80 and a FA Rate ≤ 0.30 for two consecutive sessions. Red line = 0. See **Figure S5**. **(F)** Probability density function for overall licking-related activity at each behavioral time point for the JAWS cohort. Vertical black line = sound cue representing trial start as the texture moves towards the whisker field. Vertical red lines = start (texture arrival at endpoint in whisker field) and end (texture departure towards starting point) of the PT window (time where mice can respond by licking). Colored boxes = 500ms grace period (licking does not trigger any outcomes). **(G-H)** Same as in **E, F** for the No Stim cohort. **(I)** JAWS cohort requires significantly more training sessions for expert discrimination compared to the FP and No Stim cohorts. **(J)** Longitudinal representation of sessions required for expert discrimination. **(K)** Comparison of Hit Rate, FA Rate, Sensitivity (d’), and Bias during the Learning and Expert phases. **(L)** Average RT is slower during photoinactivation than non-photoinactivated trials. Data are mean ± SEM. * p < 0.05.

## Discussion

### POm Involvement in Sensorimotor Processing

POm has received considerable attention for its potential roles in sensory/nociceptive processing,^7,12,13,14,15,16,17,43^ pain signaling,^19,20,21,126,127^ and cortical plasticity mediation.^22,23,24,25,26,^ _34,50,51_ Its widespread connectivity with sensory and motor cortical areas (S1, S2, and M1),^2,48,49,54,55,128,129^ along with its strong modulation with behavioral state and arousal,^17,18^ suggests a significant, yet still undetermined role in sensory-guided behaviors. VPM thalamus, by contrast, projects to much more delimited domains of S1, with axons coursing through striatum without forming synapses^73^. POm projections to the dorsal striatum arise from branches of the main axons bound for the cortex and terminate in a region of the dorsal striatum that overlaps with corticostriatal inputs from S1 and M1,^2,29,31,32,33,49,73,130^ potentially placing POm projections in a powerful position to influence sensorimotor integration. We aimed to elucidate the functional innervation pattern of POm onto striatal cell types, and the role of POm-striatal projections on behavioral performance and learning. Slice electrophysiology revealed strong POm-mediated synaptic inputs to D1-SPNs, D2-SPNs, and PV interneurons, with shorter latency responses in PVs. *In vivo* photometry recordings showed increasing activation of POm-striatal axons across task learning, with axonal signals and pupil dilation in expert mice increasing prior to and becoming stereotyped to stimulus presentation, independent of stimulus type or trial outcome. Photoinactivation of POm-striatal axons resulted in prolonged RTs and delayed learning, with more training sessions required to achieve expert behavioral performance. We discuss below the complex feedforward and feedback circuitry that POm is part of, and potential functional differences in POm-striatal projections compared to POm-cortical projections in sensorimotor behavior.^18,26,51^ We propose that POm could provide an early salience- or arousal-related “priming” signal to the pDLS that interacts with subsequent corticostriatal signals, leading up to behavioral action selection.

### Does POm Provide a “Priming” Signal to Striatum?

A striking feature of our POm-striatal projection measures in behaving mice was the early task-related activation, with the strongest increases before and during texture presentation rather than during trial outcome, reward presentation, or licking. SPNs require an up-state transition from their relatively hyperpolarized resting potentials to reach action potential threshold.^131^ While this up-state transition is correlated with cortical oscillatory activity,^87,132^ the role of thalamostriatal projections has not been fully elucidated.^133^ POm inputs likely arrive in striatum with shorter latency than cortical inputs following whisker stimulation,^55,56,105,134^ and therefore could be involved in initiating the up-state transition in the SPNs they innervate. We refer to this early POm-striatal input as a putative “priming” signal because it could serve to sensitize striatal neurons to respond to subsequent synaptic input. We found that POm provides large amplitude PSPs to D1- and D2-SPNs, of similar amplitude to previously measured M1 inputs and larger amplitude than S1 inputs^89^. This equally strong innervation of D1- and D2-SPN is consistent with previous work on the co-activation of SPN populations during natural movements.^135,136,137^ The early timing, together with the equal large-amplitude innervation of both direct and indirect pathways (D1- and D2-SPNs, and PV interneurons), suggest that POm projections are well-positioned to “prime” dorsal striatal circuitry for processing subsequent synaptic input.

### Impact of PV-interneuron-mediated feedforward GABAergic signaling

POm innervation of PV interneurons was particularly prominent, providing equal amplitude PSPs with shorter latency than SPNs (Fig. 1). Striatal PV cells are thought to provide feedforward inhibition to surrounding neurons via short-latency GABAergic synaptic signals.^67^ How could PV cells be involved in “priming”? First, PV cells do not exert an inhibitory effect, as defined by postsynaptic hyperpolarization, under all conditions. Stimulation of a presynaptic PV cell in synaptically coupled PV-SPN paired recordings causes GABA-mediated depolarization in the postsynaptic SPN when the SPN’s resting potentials was negative.^178^ Indeed, recent work found that the chloride equilibrium potential is surprisingly positive in adult striatal SPNs, closer to -60 mV compared to the more typical -80 mV.^138^ An approximately -60 mV equilibrium potential means that a GABA receptor-mediated chloride current produces depolarization instead of hyperpolarization when the postsynaptic neuron is at a down-state resting potential of approximately -80 mV.^138,178,179^ In Day et al. (2024), when GABA-mediated depolarization was paired with further glutamate-evoked depolarization, the probability of action potential firing was increased, suggesting that GABA-mediated depolarization was excitatory rather than shunting.^138^ In this scenario, if SPNs are sitting at a relatively hyperpolarized (down state) resting potential when POm becomes active, the disynaptic GABAergic input provided by PV interneurons could actually have a depolarizing effect on SPNs and combine with the direct glutamate-mediated depolarization provided by POm. Thus, under certain conditions, PV interneurons could contribute to “priming” striatal circuitry by adding to the early depolarization of SPNs.

The situation is different when the postsynaptic neuron is already depolarized above -60 mV (as in an up state). In this scenario, the effect of GABAergic input is hyperpolarizing, and PV cell-mediated feedforward signals to SPNs would be inhibitory, in line with traditional views.^63,65,66^ In this case, instead of enhancing the initial depolarization of striatal neurons (as above), POm input would likely provide a temporally restricted depolarizing signal, followed by hyperpolarization. POm-mediated feedforward inhibition could, depending on the intricacies of local connectivity, act to prevent up-state transitions of specific SPN ensembles, effectively increasing the signal-to-noise of striatal population activity and therefore the recruitment of SPNs by subsequent cortical or thalamic inputs. Thus, it is possible that POm projections have dual effects on striatal circuits depending on ongoing striatal activity. GABAergic signaling, mediated by PV or other types of striatal interneurons, is likely critical in mediating such state-dependent effects on postsynaptic striatal circuitry.

### POm Involvement in Salience Networks and Behavioral State

Many extrinsic and intrinsic factors influence striatal function. For example, the striatum receives inputs from myriad subcortical regions including other thalamic nuclei^73,99^ and external globus pallidus.^139,140^ A crucial factor in POm signal timing is its inhibitory gating by two GABAergic inputs, ventral zona incerta (vZI)^16,35,36,37,42^ and anterior pretectal (APT).^16,41,143^ All three nuclei receive ascending spinal trigeminal (whisker-related sensory) inputs, but vZI and APT efficiently shunt incoming sensory information via feedforward inhibition onto POm neurons.^16,39,44^ This inhibitory gating is overcome by (1) arousal-related cholinergic suppression of presynaptic GABA release within POm^43^ or (2) convergence of bottom-up and top-down signals within a specified temporal window.^31,144^ Both factors are likely at play during sensory-guided behavior. The involvement of cholinergic signaling is consistent with the POm-striatal “priming” concept if cholinergic signaling prompts POm before a behaviorally relevant period of sensation, or immediately prior to sensory information becoming perceptible.^145^ In addition to cholinergic, cortical, and subcortical GABAergic afferents, POm also receives direct glutamatergic input from superior colliculus (SC) that enhances sensory responses.^27,34^ SC, a region implicated in attentional orienting of either somatosensory^34,146^ or visually-relevant^147,148^ stimuli, bidirectionally modulates POm, further ascribing a potential arousal-related functional role. It is yet to be resolved to what extent POm-striatal inputs are driven by ascending (feedforward) sensory information, descending cortical-POm feedback, or the convergence of both, but the interplay and integration of ascending and descending signals is likely to be an essential feature in POm activation and its involvement in behavioral salience.

It is difficult to differentiate movement- and arousal-related brain states. POm neuronal activity shows a close relationship with spontaneous (task-free) whisker movements, and pupil-indexed arousal in head-restrained mice, ^18^ while in freely moving mice both VPM and POm activity correlate with head and whisker movements, ^174^ indicating that POm is generally coactive with exploratory head and whisker movements and behavioral arousal. During task performance, the situation may change with training and attentional effects. For 11xamplee, Petty and Bruno (2024) ^175^ showed that POm activity correlates more closely with task demands than tactile or visual stimulus modality. Our data indicate that POm-striatal axonal signals are increased at trial start during anticipation of tactile stimulus delivery and through the sensory discrimination period, then decrease to baseline levels during licking and water reward collection (Fig. 3). Results of ^175^ together with ours suggest that POm is particularly active during the context of behaviorally relevant task performance. Thus, we think that pupil dilation indexes general movement and arousal, which POm correlates with during spontaneous behaviors, but that POm activity becomes more specific and more prominent during the movement, anticipation, and arousal associated with learned sensorimotor behavioral tasks.

### Parallel Thalamostriatal Pathways, Additional Striatal Cell Types

Pf is the best studied thalamic nucleus projecting to the striatum. A major difference between POm and Pf striatal innervation is Pf innervation of cholinergic (ChAT; tonically active) interneurons.^72,87,90,102,104,149,150^ We did not determine whether POm innervates ChAT interneurons, but available evidence suggests it might only weakly. Retrograde tracing to map brain-wide inputs to striatal ChAT interneurons showed that Pf provides the majority of thalamic innervation of striatal ChAT neurons compared to other thalamic nuclei.^149^ Many other thalamic nuclei, including POm, showed little or no labeling. However, it is possible that these thalamic nuclei, including POm, provide functional innervation of ChAT interneurons that is insufficiently assessed by anatomical methods alone.

Another difference between Pf and other thalamic nuclei innervation of striatum is the subcellular localization of their synaptic inputs. Pf synapses tend to localize to dendritic shafts of SPNs rather than dendritic spines^151,152,153^ whereas other thalamic nuclei form synapses onto SPN dendritic spines.^176^ These differences could affect postsynaptic signal processing, such that Pf shaft inputs may induce a longer temporal window for integration via dendritic filtering,^154,155^ whereas POm spine inputs may cause a more restricted temporal window for signal integration. These properties could influence the behaviorally relevant time window of thalamostriatal and corticostriatal signal integration^86,89,90^, for example, with POm requiring time-locked inputs onto the same or nearby spines for maximal postsynaptic effect.^56^ The occurrence and relevance of such mechanisms in different thalamostriatal pathways will require further investigation. Additionally, it will be important to understand the innervation patterns of POm-striatal projections beyond the three cell types we have studied here, including other interneurons responsive to thalamic stimulation such as tyrosine-hydroxylase (THIN) and neurogliaform (NGF) interneurons.^67,109^ Lastly, the effects of dopamine modulation of thalamostriatal signaling directly^62^ and indirectly via cholinergic interneurons^141,142^ would be an important area of further study.

### POm as a Link between Cortical and Striatal Signaling

POm axons bifurcate and contact neurons within pDLS,^49,52,53,54,55,57,58^ but the main terminations continue to cortex where they strongly innervate L1 and L5a of S1.^48,49,54,55^ POm input excites L5 pyramidal neurons (and L2/3 neurons) via L5 perisomatic synapses as well as distal dendritic synapses in L1.^50^ Synaptic plasticity of POm-cortical inputs and increased excitation of L2/3 and L5 neurons has been implicated in sensory learning.^22,23^ Given the capacity of POm-cortical synapses for learning-related plasticity, it will be important to determine whether POm-striatal synapses also undergo learning-related plasticity and how it may be distinct from POm-cortical plasticity. Remarkably, POm axonal branches, even from the same neuron, display synaptic terminals with distinct structure and functional properties in M1 compared to S1 ^177^; thus it is possible that the striatal synapses of POm neurons could undergo distinct changes with learning compared to their cortical counterparts.

How the divergent cortical and thalamic projections of POm neurons may be involved in coordinating thalamostriatal and corticostriatal signals is another important issue. Striatal neurons that receive POm input are likely also to receive cortical L5a input, because L5a neurons comprise the predominant S1-striatal projection.^68,69,76^ Thus a disynaptic POm-S1-pDLS loop exists in addition to the monosynaptic POm-pDLS projection. During behavior, striatal circuitry may be initially engaged via the monosynaptic POm projection followed, in tens of milliseconds, by the disynaptic loop that recruits L5a cortical pyramidal neurons to provide input to already depolarized (“primed”) striatal neurons. On the hundreds-of-milliseconds time scale of a behavioral trial, it is possible that these pathways are active multiple times, with L5b cortical feedback activating POm _31,33_ after the initial feedforward volley of activity, initiating POm-driven signaling again. It is possible that an individual SPN receives both POm and L5a input, with the POm input providing the initial “priming” depolarization, followed by further depolarizing input from a POm-driven L5a neuron. It is unclear whether thalamic “priming” of striatum is necessary for corticostriatal activation of SPNs. Based on the recurrence and convergence of these pathways, it is possible that POm acts to synchronize thalamo- and corticostriatal signals under certain behaviorally relevant conditions. This would be a unique role for POm compared to Pf, since Pf strongly innervates striatum but only weakly innervates cortex.

### POm-striatal Recruitment during Learning: Unresolved Mechanisms

Behavioral state^18^ influences the efficiency of sensorimotor learning.^26,51^ Overall, we observed a learning-related increase in POm-striatal axonal activity (as measured by photometry) that correlated with pupil dilation in many but not all phases of behavioral performance, and a necessity of these projections for efficient behavioral performance and learning, supporting a role in task-related behavioral arousal. Photometry signals and pupil dilation in expert mice were tightly correlated between trial onset and texture presentation, but decoupled when photometry signals decreased prior to licking and reward delivery. Sorting trials by presented texture (stimulus) or trial outcome (response), photometry signals remained elevated, while both licking and pupil dilation exhibited stimulus- or response-specific changes. This suggests that POm-striatal projections do not encode specific sensory- or outcome-related information, but rather arousal or salience during anticipatory states of a learned behavior. The effects of photoinactivation of PO-striatal projections included delayed RTs on individual trials and more sessions required to achieve expert performance. These effects suggest that POm serves to enhance striatal signaling during increased behavioral states associated with performance of learned sensorimotor behavior. POm activation early during behavioral trials may set the stage for other important inputs, such as corticostriatal inputs, to influence action selection. The increased amplitude of POm-striatal signals with learning suggests an increased influence of POm on striatum, and thereby on goal-driven sensorimotor behaviors. POm likely becomes increasingly tuned to task-specific behavioral arousal and anticipation than to general movement-related arousal.

An important unresolved issue is how POm-striatal signals increase with learning. Inherent in our measures of POm-striatal axons is the limitation that photometry only captures a bulk axonal signal and lacks the ability to resolve individual axons. Therefore, we were unable to resolve whether the increased bulk signal that we observed was due to a uniform increase across all POm-striatal axons or a heterogeneous increase present only in specific axons. This is important because POm neurons exhibit heterogeneous responses to direct paralemniscal stimulation,^10^ peripheral stimulation,^15^ and the suppression of vZI activity,^39^ and also show functionally distinct anterior and posterior subpopulations.^2,49,54,130,156^ Therefore, we might expect heterogeneous learning-related changes in activity within the POm neuronal population, a mechanistic question that would be important to address in future studies.

Although we placed the optical fiber above pDLS to specifically record from axons, rather than somas, it is possible that POm-cortical axons of passage contributed to the recorded signal. However, even if signal contamination were present, we think the measures would be largely similar, as most POm-cortical projections bifurcate within striatum rather than projecting exclusively to either striatum or cortex.^49,55^ Signal modification is more likely to occur at synapses via presynaptic or postsynaptic mechanisms than at the main axon.^157,158,159^ In future work it would be valuable to investigate the intriguing possibility that POm projections have discrete target-specific functions, such that POm-striatal signals may be involved in distinct aspects of sensorimotor behavior compared with POm-cortical signals.^18,26,51^

In summary, we show that POm-striatal projections encode a behaviorally relevant arousal-related signal that increases with learning. POm-striatal projections may constitute a “priming” signal that combines with corticostriatal signals and contributes to inducing the up-state transition of SPN ensembles involved in action selection, by equally engaging D1- and D2-SPNs and PV interneurons. These finding suggest a previously unknown functional role of POm engagement of striatal microcircuitry. It will be important for future studies to investigate whether POm further innervates other striatal interneurons, to assess the timing between POm-striatal and corticostriatal inputs to SPNs, and to assess whether POm-striatal synapses undergo synaptic plasticity across learning.

## Acknowledgements

The authors would like to thank all members of the Margolis lab for their comments related to manuscript text and figures. This work was generously supported by funding from the National Institutes of Health (F31NS117093, A.J.Y.); (NCATS TL1TR003019, A.J.Y.); (R01NS094450, D.J.M.), the National Science Foundation (IOS-1845355, D.J.M.), and the Rutgers Busch Biomedical Grant Program (I. L./D.J.M.).

## Author Contributions

A.J. Yonk and D.J. Margolis conceived and designed the research strategy. A.J. Yonk performed and analyzed electrophysiology experiments with assistance from M.A. Gradwell and A.J. George. A.J. Yonk performed and analyzed fiber photometry behavioral experiments with assistance from I. Linares-Garcia, L. Pasternak, S.E. Juliani, and A.J. George. A.J. Yonk performed and analyzed optogenetic behavioral experiments with assistance from I. Linares-Garcia and A.J. George. A.J. Yonk prepared the figures with input from I. Linares-Garcia, S.E. Juliani, M.A. Gradwell, A.J. George, and D.J. Margolis. A.J. Yonk wrote the paper, D.J. Margolis revised the paper after review, with all authors contributing to its editing. A.J. Yonk and D.J. Margolis acquired funding. D.J. Margolis supervised the study. All authors approved the final version of the manuscript.

## Methods

### Data and Code Availability

Data are available upon reasonable request from the lead contact, David Margolis and at github.com/margolislab.

### Animals

All work involving animals including housing, surgery, behavioral experimentation, and euthanasia was approved by the Rutgers Institutional Animal Care and Use Committee (protocol #: 999900197). Mice were group housed in a reverse light cycle room (lights on from 20:00 to 08:00) with food and water available *ad libitum* with the exception of mice undergoing water restriction during behavioral experiments. All handling and experiments were conducted within the dark phase of this light cycle. Regardless of their experimental designation, all experimental animals underwent a unilateral AAV injection or a simultaneous AAV injection and cannula implant between –5 - 65 days (average: 52.37 days, range: –7 - 61 days). Briefly, male and female double transgenic mice were used for electrophysiology experiments. To identify specific neuronal populations during electrophysiology, Ai14 mice (cre-dependent tdTomato; The Jackson Laboratory, #007914) were crossed with either (1) D1-SPN cre (B6.FVB(Cg)-Tg(DrD1cre)EY262Gsat/Mmucd; MMRRC, #030989), (2) D2-SPN cre (B6.FVB(Cg)-Tg(Adora2a-cre)KG139Gsat/Mmucd; MMRRC, #036158), or (3) PV-cre (B6.129P2-Pvalb^tm1(cre)Arbr^/J; The Jackson Laboratory, #017320) mice. This permitted red fluorescence in the specific cell population for visual identification via epifluorescent illumination. Animals designated for electrophysiology were euthanized between–2 - 4.5 months. Both male and female wild-type mice (The Jackson Laboratory #000664) were used for fiber photometry and optogenetic experiments. To motivate behavioral performance, daily water intake was restricted to ∼1.5mL per mouse per day. Body weight was carefully controlled and never permitted to drop below 80% of a calculated baseline value.^169^ The number of male and female mice were as follows, by experiment type: 6 male, 4 female (electrophysiology); 3 male, 2 female (fiber photometry); 4 male, 5 female (optogenetics). Data were not analyzed for sex differences.

### Electrophysiology and Adeno-Associated Viral (AAV) Injection

Male and female double transgenic mice designated for electrophysiology experiments underwent a unilateral injection of channelrhodopsin-2 (ChR2; pAAV-hSyn-hChR2(H134R)-EYFP; Addgene #26973)^160^ targeting the left posterior medial (POm) thalamic nucleus. Briefly, mice were anesthetized using isoflurane (4% induction, 1-2% maintenance) and moved into a stereotaxic apparatus (Stoelting/Kopf Instruments) containing a feedback-controlled heating pad on the base (maintained between 35-37°C; FHC). Ophthalmic ointment (Akorn) was applied to the eyes to prevent them from drying out. Ethiqa XR (3.25 mg/kg; Fidelis Animal Health) and Bupivacaine (0.25%, 0.1mL, Fresenius Kabi) were injected subcutaneously into the right flank and scalp, respectively. After, the scalp was sterilized by three cycles of Betadine (Purdue Products) and 70% ethanol, a midline incision was made. The exposed skull was cleared of fascia and leveled by confirming that bregma and lambda coordinates were on the same dorsoventral plane. A craniotomy was drilled (coordinates with respect to bregma: anteroposterior (AP) = -2.05 mm, mediolateral (ML) = +1.35 mm, dorsoventral (DV) = -3.25 mm) and the micropipette containing ChR2 was slowly lowered to the appropriate depth and permitted to sit for 5 minutes. Following this, ∼100 nL of ChR2 viral solution was injected over the course of ∼15 minutes via the Nanoject III system (Drummond Scientific). After an additional delay of 12 minutes, the micropipette was slowly raised, the scalp was sutured and secured with tissue glue. Immediately following surgery, mice were transferred to a clean cage on top of a heating blanket until ambulation was observed. Mice were continually monitored for 72 hours post-surgery. After this observation period, mice were permitted to recover for at least three weeks before undergoing electrophysiological experiments, permitting ChR2 expression into POm axon terminals in the striatum.

### Whole Cell Patch Clamp Recordings

Mice were briefly induced (via 3% isoflurane), deeply anesthetized with an intraperitoneal injection of ketamine-xylazine (300/30 mg/kg, respectively), and transcardially perfused with recovery artificial cerebrospinal fluid (ACSF), which contains the following (in mM): NMDG 103, KCl 2.5, NaH_2_PO_4_ 1.2, NaHCO_3_ 30, HEPES 20, Glucose 25, HCl (1N) 101, MgSO_4_ 10, Thiourea 2, Sodium Pyruvate 3, N-Acetyl-L-Cysteine 12, and CaCl_2_ 0.5 (saturated with 95% O_2_ and 5% CO_2_).^89,109^ Following decapitation, the brain was rapidly extracted and submerged in recovery ACSF before being mounted onto a VT-1200S vibratome (Leica). The vibratome chamber was filled with oxygenated recovery ACSF, and 300µm slices were cut. Slices were immediately transferred to recovery ACSF that was heated to 35°C for ∼5 minutes. After, slices were transferred to RT external ACSF which contained (in mM): NaCl 124, KCl 2.5, NaHCO_3_ 26, NaH_2_PO_4_ 1.2, Glucose 10, Sodium Pyruvate 3, MgCl_2_ 1, and CaCl_2_ 2 (saturated with 95% O_2_ and 5% CO_2_), and slices were allowed to recover for at least ∼45 minutes before recording. Once the hippocampus began to appear, vibratome sectioning was terminated, and the posterior tissue block containing the injection site was transferred into 10% neutral-buffered formalin for post-hoc confirmation.

Whole-cell patch clamp recordings were acquired from slices that were constantly superfused (2-4mL/min) with oxygenated external ACSF at ∼34°C. Slices and cells were visualized by infrared differential interference contrast (IR-DIC) microscopy using a CCD camera (Hamamatsu) mounted onto a BX-51WI upright microscope (Olympus) fitted with a swinging objective holder containing two switchable lenses: a 4X air lens and a 40X water-immersion lens. Patch pipettes (3-5 MΩ) were pulled from borosilicate glass micropipettes (2mm O.D., Warner Instruments) using a P-1000 horizontal puller (Sutter Instruments).

Current-clamp recordings were obtained from unidentified and identified striatal neurons in mice expressing tdTomato in either D1-SPNs, D2-SPNs, or PV cells within pDLS (-0.34 to - 1.22mm relative to bregma), which is known to receive POm projections.^55,56,105^ The internal pipette solution for current-clamp experiments contained (in mM): K Methanesulfonate 130, KCl 10, HEPES 10, MgCl_2_ 2, Na_2_ATP 4, Na_2_GTP 0.4, at pH 7.25 and 285-290 mOsm/L. Further, 2% biocytin was freshly dissolved in the internal solution on the recording day. ChR2 in the POm axon terminals was stimulated via illumination with a 2.5 ms, 470 nm LED light pulse (∼0.6 mW measured after the objective; Thorlabs) delivered through the 40X objective lens. The illumination spot size had a diameter of ∼550 µm. After patching, the internal solution was permitted to dialyze for ∼5 minutes. At this point, the resting membrane potential was recorded. All cells were held at -80 ± 2mV to ensure equal driving forces when studying synaptic strength and short-term synaptic plasticity. Baseline voltages that drifted outside of this range were excluded from analysis. Patched cells were held for ∼25-35 minutes while being run through a standardized set of protocols: (1) hyperpolarizing/depolarizing current steps, (2) single pulse (SP), (3) paired pulse ratio (PPR), and (4) train stimulation. Occasionally, unidentified cells within the same FOV were sequentially patched following the initial identified cell patch to control for injection site volume and location. After breaking in, the cell was allowed to recover for ∼5 minutes before being subjected to hyperpolarizing and depolarizing current steps (-300 pA to +400 pA, 50 pA steps, 500 ms, 15 sweeps) for cell health and intrinsic parameter confirmation. For the SP protocol, a single 2.5 ms blue light pulse was presented once every 15s for 20 sweeps. For the PPR protocol, five 2.5 ms pulses, separated by 50 ms inter-pulse intervals (IPI), were presented once every 15s for 20 sweeps. For the train protocol, thirty 2.5 ms pulses, separated by 64.2 ms IPI (15 Hz), were presented once every 30s for 5 sweeps. While these protocols were being run, biocytin within the internal solution diffused into the cell for subsequent 3D morphological reconstructions. Data were acquired via a EPC10USB amplifier and digitized at 20 kHz in Patchmaster Next (HEKA). Liquid junction potential was not corrected in these traces.

### Analysis of Patch Clamp Recordings

All data were analyzed offline using custom-written MATLAB and Python scripts. DAT files from Patchmaster Next (HEKA) were imported, organized, and saved as a mat variable. The mat variable data was imported into Python for post-processing using the electrophysiology feature extraction librarIFEL) created by the Blue Brain Project.^161^ All pertinent intrinsic value parameters were calculated at the +350pA current step. Key values pertaining to every action potential in a sweep were calculatIia eFEL including the action potential threshold value and index, the peak value and index, and the corresponding minimum afterhyperpolarization (AHP) value and index. Each action potential threshold value and index were captured using a derivative threshold method (dV/dt ≥ 10mV/s). Action potential properties were assessed for all spikes in a sweep and averaged together. The action potential peak was defined as the difference between the action potential threshold and its maximum positive peak. Half-height width (HHW) and rise time were both calculated via interpolation. HHW was measured at 50% of the action potential peak, while rise time was measured from 10-90% of the action potential peak. AHP amplitude was calculated as the difference between the action potential threshold value and the minimum AHP value. Maximum frequency

For all optogenetically-evoked postsynaptic potentials (PSPs), a baseline measure was calculated by averaging the first 10,000 sampling points for each individual sweep. This measured value was subtracted from every value in each individual sweep, setting the baseline equal to zero. The maximum PSP amplitude, relative to the zero baseline value, was calculated from the average of 20 sweeps during SP and PPR protocols, and 5 sweeps during Train protocols. The latency to maximum PSP amplitude was measured as the difference between photostimulation onset and the maximum PSP index. For PPR and Train protocols, an exponential function was fitted to the decay component of each PSP, and their amplitudes were extracted after subtracting the decay of preceding PSPs.

### Simultaneous AAV Injection and Optical Cannula Implantation Surgery

For male and female wild-type in the designated fiber photometry cohort, a 400µm core optical cannula (ferrule OD = 2.5mm, length = 2mm, nA = 0.50; Thorlabs #CFM15L02) was implanted directly above the left pDLS along with a ∼100nL unilateral injection of axon-jGCaMP8s (pAAV-hSynapsin-axon-jGCaMP8s-P2A-mRuby3; Addgene #172921)^162^ into the ipsilateral POm. Male and female wild-type mice designated for optogenetic manipulation were split into two cohorts: (1) the No Stim or (2) the JAWS cohort. Both optogenetic cohorts were implanted with a 200µm core optical cannula (ferrule OD = 1.25mm, length = 2mm, nA = 0.50, CFMLC52L02, Thorlabs) above the left pDLS along with a ∼100nL AAV injection into the ipsilateral POm. However, the No Stim cohort was injected with the excitatory optogenetic actuator, ChR2 (pAAV-hSyn-hChR2(H134R)-EYFP, Addgene #26973),^160^ while the JAWS cohort was injected with the inhibitory optogenetic actuator, JAWS (pAAV-hSyn-JAWS-KGC-GFP-ER2, Addgene #65014).^123,124,125,163^ Further, an HHMI head plate^164^ was fitted to each mouse using methods previously described.^89,117,165,166,167^ Briefly, mice were anesthetized with isoflurane (4% induction, 0.8 – 1.5% maintenance) and mounted within a stereotaxic frame (Stoelting/Kopf Instruments) with a feedback-controlled heating pad (FHC) maintaining the body temperature between 35 – 37°C. Following administration of an analgesic (Ethiqa XR, 3.25mg/kg; Fidelis Animal Health) to the left flank and a local anesthetic (0.25% Bupivacaine; Fresenius Kabi) under the scalp, the scalp was sterilized with a triple cycle of Betadine (Purdue Products) followed by 70% ethanol (Fisher). A midline incision was made, and a circular piece of scalp was removed to expose the skull. Both lateral muscles and the nuchal muscle were separated from the skull. The skull was cleaned by gently scraping away the periosteum. A dental etch bonding agent (iBond; Heraeus Kulzer) was applied to the clean skull and cured with blue light for 20s twice. A ring of dental composite (Charisma; Heraeus Kulzer) was applied to the outer edge of the skull and cured with blue light. After ensuring that bregma and lambda coordinates were equal in the dorsoventral plane, two craniotomies were made: one above the left posterior striatum (AP = -0.80mm, ML = +2.8mm, DV = 1.8mm) for cannula implantation and the other above left POm (AP = -2.05mm, ML = +1.35mm, DV = -3.25mm) for the corresponding AAV injection. The AAV injection was always performed first. The micropipette containing axon-jGCaMP8s (fiber photometry cohort), ChR2 (No Stim cohort), or JAWS (JAWS cohort) was slowly lowered to the appropriate depth and permitted to sit for 5 minutes. ∼100nL of AAV solution was injected over the course of ∼15 minutes via the Nanoject III system (Drummond Scientific). After an additional delay of 12 minutes, the micropipette was slowly raised. Once the optic cannula was secured in the stereotaxic frame and lowered to the appropriate DV coordinate, it was secured with dental composite (Tetric Evoflow; Heraeus Kulzer) that was cured with blue light. The HHMI headpost was delicately secured on the posterior area of the charisma ring with cyanoacrylate and dental composite. Finally, a single layer of dental composite was used to build and secure the rest of the headcap before being cured four times with blue light for 20s each. The scalp was closed around the headcap by using cyanoacrylate.

Immediately after the optical implant and viral injection surgery, mice were placed in a sterile, clean cage that was half-on/half-off a heating pad until ambulation was observed. Mice were diligently monitored for 72 hours post-surgery. After this monitoring period, mice were transferred to a clean cage on a ventilated rack for at least three weeks prior to handling and water restriction.

### Handling and Water Restriction

After a three-week waiting period, mice were placed under citric acid water restriction^168^ during which mice were handled twice daily for one week. The bitterness of citric acid naturally causes mice to reduce their water consumption and, consequently, their weight while still having access to water. Initially, mice were acclimated to handling as researchers placed their hands into the cage for increasing amounts of time (e.g., 5 minutes to 10 minutes to 15 minutes). After mice became comfortable, they were held for increasing amounts of time (e.g., 2 minutes to 5 minutes to 10 minutes). Additionally, mice were acclimated to head fixation by holding their HHMI headposts (e.g., 30 seconds to 1 minute to 2 minutes), and they were allowed to freely explore the behavioral tube for 5 minutes per handling session. Finally, mice were headfixed for increasing amounts of time in the behavioral setup (e.g., 5 minutes to 10 minutes to 15 minutes). Water was provided via transfer pipette to comfortably acclimate mice to head fixation. The head-fix apparatus contained a tube (length = 15cm; inner diameter = 4cm) that was affixed to a custom metal platform (length = 17cm; width = 12cm). The platform also contained HHMI mounting arms and holders for head fixation.^164^ During the final day of handling, mice were transitioned to full water restriction as this permits greater control over motivation level. During behavioral testing, daily water intake was limited to ∼1.5mL per day to motivate performance on the Go/NoGo paradigm described below. The baseline body weight was measured daily during water restriction, and overall body weight was not permitted to drop below 80% of the baseline weight, consistent with levels of restriction used to motivate performance.^169^

### Fiber Photometry Setup

Fiber photometry data were collected using a RZ10x lock-in amplifier within the Synapse suite (Tucker-Davis Technologies). This amplifier and accompanying Synapse software was used to control a custom-built optical benchtop through drivers (LEDD1B, Thorlabs) to modulate LED signals. Briefly, this optical benchtop consisted of a self-contained system of four 30mm cage cubes with integrated filter mounts (CM1-DCH/M, Thorlabs). A 405nm LED (M405L4, Thorlabs) and a 470nm LED (M470L3, Thorlabs) were mounted onto the first 30mm cage cube. The 405nm LED was used to extract the calcium-independent isosbestic signal, and the 470nm LED was used to acquire calcium-dependent axonal GCaMP signals during the Go/NoGo paradigm. The 405nm isosbestic signal was modulated at 210Hz, and the 470nm GCaMP signal was modulated at 330Hz. A 425nm dichroic longpass filter (DMLP425R, Thorlabs) in the first cage cube reflected the 405nm excitation light and permitted the 470nm light to pass through. As the light entered the second cage cube, both excitation lights were reflected by a 495nm dichroic longpass filter (495DCLP, 67-079, Edmund Optics) into the third cage cube. A 460/545nm bandpass filter (69013xv2, Chroma) reflected both excitation wavelengths down to the subject via a low autofluorescence patch cable (MAF3L1, core = 400µm, nA = 0.50, length = 1m, Thorlabs). This cable was attached onto the implanted optical cannula (see above) by a ceramic mating sleeve (Thorlabs). Isosbestic and axon-jGCaMP8s emissions were collected via the optic cannula and passed through the 460/545nm bandpass filter (69013xv2, Chroma)into the fourth. Finally, the emission fluorescence passed through the detection pathway to reach the RZ10x photosensors for online observation.

### Orofacial Video Recording

POm activity has been well correlated with whisking and pupil activity.^18,26^ To analyze these dynamics, an LED driver controlled an IR spotlight that illuminated the contralateral eye and mystacial pad, and facial recordings were captured through an autofocusing USB webcam (NexiGo N660P) at ∼20fps within the Synapse software. To limit light pollution from outside sources (e.g., LED illumination within the brain/eye), an IR filter was placed in front of the webcam. Also, a shortpass emission filter at 750 nm (Chroma #ET750sp-2p8) was placed between the third and fourth cage cubes to prevenI recorded IR light from overloading the photosensors.

### Synchronization of Task-Related Components

The Synapse software suite (Tucker-Davis Technologies) permitted the synchronous recording of both emitted (isosbestic and calcium) signals and video frames. Furthermore, the LabVIEW system-controlled paradigm-related components (e.g., texture movement, lick thresholds, trial type, trial outcome, etc.) and recorded the resulting behavioral parameters (e.g., licking activity, trial type, and trial outcome). To synchronize these two data streams for post-hoc analysis, TTL pulses relating to the texture arrival at its endpoint (which indicates the start of the presentation time window) from the LabVIEW system were captured within the Synapse software.

### *In Vivo* Optogenetics

For the JAWS cohort, a high-powered 617nm LED (M617F2, Thorlabs) and current driver (LEDD1B, Thorlabs) were used for photoinactivation of POm axons in the striatum.^123,124,125,163^ LED stimulation was provided at a probability of 0.50 for every session from the start of the Learning phase until mice reached the Expert phase.^26^ The light intensity was measured at ∼7mW at the tip of the fiber. Stimulation intensity was kept consistent between mice and days by measuring the intensity with an optical power meter (PMD-100D, Thorlabs) prior to the first session every day. Light was delivered to the thalamic afferents in pDLS through an optical fiber patchcord (M95L01, fiber core diameter: 200µm, length: 1m, nA: 0.50) connected to a 200µm core optical cannula (described above) via a mating sleeve (Thorlabs). A small piece of black heat shrink tubing (Qualtek) was placed over the cannula during LED testing to prevent stray light from illuminating the presented texture during the task. The LabVIEW system controlled a Pulse Pal^170^ that activated the LED current driver to provide constant illumination for two seconds, evenly split 1s before and 1s after texture presentation, corresponding to the increased calcium activity observed during fiber photometry recordings.

For the No Stim cohort, a high-powered 470nm LED (Prizmatix) and current driver (Prizmatix) were used for optogenetic activation. Note that no LED stimulation was provided during the Learning or Expert phase. Testing occurred during the first 4 sessions after the 5 initial Shaping sessions and the last 4 sessions after mice reached Expert status. Sessions consisted of 50 baseline trials followed by 10 alternating blocks of 10 OFF and 10 ON trials. The light intensity was measured at ∼5mW at the tip of the fiber and was kept consistent between mice and days by measuring the intensity with an optical power meter (PMD-100D, Thorlabs) prior to the first session on testing days. Light was delivered to the thalamic afferents in pDLS through an optical fiber patchcord (M73L01, fiber core diameter: 200µm, length: 1m, nA: 0.50, Thorlabs) connected to a 200µm optical cannula (described above) via a mating sleeve (Thorlabs). A small piece of black heat shrink tubing (Qualtek) was placed over the cannula during LED testing to prevent stray light from illuminating the presented texture during the task. The LabVIEW system controlled a Pulse Pal^170^ that activated the LED current driver to provide pulsed illumination for 2s at 15Hz (matching the electrophysiology train photostimulation paradigm), evenly split 1s before and 1s after texture presentation.

### Go/NoGo Whisker-Based Discrimination Paradigm

Headfixed, water restricted mice were trained to perform a whisker-based discrimination paradigm as two textures were presented unilaterally to the right whiskers in a randomized order based on a custom-written LabVIEW code (National instruments). This code used transistor-transistor logic (TTL) pulses to control all aspects of the paradigm including a water delivery spout connected to a piezo film sensor (MSP1006-ND; Measurement Specialties), and a motorized linear stage (T-LSM100A; Zaber) with a stepper motor (T-NM17A04; Zaber) containing two windmill arms holding two different sandpaper textures (Go texture = 100 grit sandpaper, P100; NoGo texture = 1200 grit sandpaper, P1200; 3M) as previously described.^89,117,165,166,167^ Mice were trained to discriminate between these two textures by licking the piezo film sensor spout. Mechanical spout displacement resulted in transient voltage changes, and a lick was defined as voltage changes crossing either an upper or lower threshold once. After a lick was detected within the appropriate response window, the LabVIEW software immediately delivered the appropriate output: for Go trials, mice received a small water reward via a solenoid valve (0127; Buerkert) through the piezo spout, and, for NoGo trials, mice received a timeout period with co-occurring white noise. This paradigm occurred within a darkened room to minimize non-tactile related cues. Both textures and the headfix apparatus were cleaned with 70% ethanol between each mouse. If mice were not performing the task, sessions could be ended early. Water could be automatically delivered (AutoReward or AR) by the experimenter following 20 consecutive trials without a response. Finally, a session could be terminated if a mouse did not lick when water was present on the end of the spout after three ARs.

Trials began with a 1000ms pre-task interval followed by a brief cue tone (100ms, 2039Hz) that accompanied windmill texture movement towards the mice. Once the windmill texture had reached a predetermined distance within the whisking field, mice had to respond within a 2000ms presentation time (PT) window. For the first 500ms of the PT window, a grace period was present where responses did not trigger appropriate outcomes to reduce impulsivity. If mice licked in response to the Go texture, the trial was considered a Hit and resulted in a water reward accompanied by a correct tone (2793Hz). If mice licked in response to the NoGo texture, the trial was considered a False Alarm (FA) and resulted in punishment parameters including a time-out period (12000ms) and an accompanying white noise during the time-out period. Air puffs were eschewed as a punishment parameter to permit continuous pupil dynamic recording. If mice did not lick to the Go or the NoGo texture, nothing occurred, and the trial was considered a Miss or a Correct Rejection (CR), respectively. Immediately after a lick was recorded or the 2000ms PT window had elapsed, the windmill texture retreated to its original position where the current texture either remained or the other texture was rotated into position. Finally, trials were separated by a 2000ms intertrial interval. Behavioral performance was tracked across texture discrimination sessions by computing multiple behaviorally-related parameters including Hit Rate, FA Rate, Sensitivity (d’) and Bias.^171^ For the fiber photometry cohort, the 405nm and 470nm LEDs provided constant illumination throughout all trials. For the JAWS cohort, LED photoinactivation occurred at a trial probability of 0.50 for every session from the start of the Learning until the end of the Expert phase.^26^ For the No Stim cohort, LED stimulation did not occur during training (i.e., during either the Learning or Expert phases). It only occurred for 4 sessions after the initial 5 shaping sessions, and 4 sessions after mice reached expert status.

This whisker-based discrimination paradigm lasted up to 3 weeks. The FP cohort were only tested once per day to limit photobleaching. The JAWS and No Stim cohorts were tested twice daily. Each session consisted of 150 trials. All cohorts progressed through three behavioral phases (Shaping, Learning, and Expert) that were segmented into five analytical time points (Shaping, Early Learning, Late Learning, and Expert). The Shaping time point was the same for all mice. For the first three sessions, mice were acclimated to reliably trigger water delivery by licking the water spout. During these sessions, neither texture was presented to the whiskers. After, mice proceeded to texture discrimination training still under the Shaping phase. For the last two sessions, both textures were presented simultaneously with a Go texture probability of 0.90 and 0.75, respectively. Go and NoGo trials were interleaved in a pseudorandom order determined by the LabVIEW software. After the 0.75 probability session, mice progressed into the Learning phase. For all following sessions, the Go texture probability was set to 0.50 with a maximum of three consecutive presentations of the same texture. Discrete behavioral time points were determined as follows. The Early Learning time point was considered the first two sessions of the Learning phase. The Late Learning time point was considered the last two sessions of the Learning phase before achieving expert status. This occurred when mice had a Hit Rate ≥ 0.80 and a FA Rate ≤ 0.30 for two consecutive sessions. A strict sensitivity threshold was not used due to artificially increased sensitivity (discrimination) values as Hit Rate and/or FA Rate approach their extremes (e.g., see FPOm-18 sessions 8 to 15 in Figure 6). After discrimination training (i.e., achieving expert status), mice in the fiber photometry cohort were subjected to a single Reward session to assess calcium, licking, and pupil activity in the absence of texture input and licking-related outcomes. During this session, the Go trial probability remained at 0.50, and the Zaber motor moved the windmill texture holder towards the whisker field, but the textures were rotated out of whisker range. Further, the upper and lower thresholds were set so that licking could not trigger outcomes before the end of the PT window. A water reward was automatically delivered at the end of the PT window during Go trials only. A whisker trim control session was performed in a subset of mice to confirm that mice used their whiskers to discriminate as previously observed.^89^

### Histology

For electrophysiology experiments, the tissue block containing the injection site was stored overnight in 10% neutral-buffered formalin at 4°C. After, it was transferred into 0.2% PBS Azide at 4°C until sectioning. The tissue block was mounted onto a stage and sectioned in 0.1M PBS at a thickness of 100µm using a VT-1000 vibratome (Leica). Slices were mounted onto microscope slides using DAPI Fluoromount-G (Southern Biotech #0100-20) and coverslipped before confocal imaging.

Following all behavioral experiments, mice were anesthetized with an intraperitoneal injection of Ketamine-Xylazine (120mg/kg Ketamine, 24 mg/kg Xylazine) and transcardially perfused with PBS followed by 10% neutral-buffered formalin. The brain was delicately extricated and stored in 10% neutral-buffered formalin overnight at 4°C. Tissue was mounted onto a stage and sectioned at 100µm using a VT-1000 vibratome (Leica). Sections were mounted and coverslipped using DAPI Fluoromount-G (Southern Biotech #0100-20). Confocal images were acquired using a LSM800 confocal laser scanning microscope (Zeiss) for injection site location verification, cannula placement, and viral expression in POm axons within pDLS and stereotypical POm-cortical projections in S1 L1 and L5a of all experimental mice.^4,25,26,47,49^ All data were acquired using the Zen software suite (Zeiss).

### Quantification and Statistical Analysis Behavioral Responding Analysis

Behavioral performance was tracked across texture discrimination sessions by computing multiple behaviorally-related parameters including Hit Rate, FA Rate, Sensitivity (d’) and Bias.^171^ Hit Rate was calculated as follows: [Hit / (Hit + Miss)], where hit is the number of correct Go trials and Miss is the number of incorrect Go trials. The FA Rate formula is similar to the Hit Rate formula except it replaces Hit with FA and Miss with CR: [FA / (FA + CR)]. Sensitivity illustrates the ability to discriminate between the signal (Go texture) and the noise (NoGo texture), and it is derived from signal detection theory.^171^ It is calculated as follows: [normalized inverse–(Hit Rate) - normalized inverse (FA Rate)]. Finally, Bias illustrates the overall responding bias, independent of trial type. It is calculated as follows: [0.5 * (normalized inverse (Hit Rate) + normalized inverse (FA Rate)].

### Pupil Analysis

After all data were captured, the recorded orofacial videos were analyzed using DeepLabCut, a deep learning model for pose estimation, to estimate pupil dynamics as mice learned to discriminate between the two textures.^118,119^ Briefly, a model^120^ was created with eight markers circumscribing the pupil, permitting the estimation of the approximate pupil area for each frame. All videos were cropped to a smaller dimension (230 x 275 pixels) that focused on each mouse’s face. For each mouse, one video was selected at varying behavioral time points, and 30 frames were extracted and manually labeled with the eight markers. Each marker corresponded to a pupil location: top, top right, right, bottom right, bottom, bottom left, left, and top left. Additionally, another marker was placed on a static location (e.g., the water spout) to label frames when blinking occurred. Overall, the initial training dataset contained 150 labeled frames, and the model was trained on this dataset with a ResNet50-based neural network for 250,000 iterations. After, the initial training videos were analyzed to assess the performance of the model from its marker estimations. 25 outlier frames from the initial training videos were extracted, manually corrected, and merged with the initial dataset. The model was trained again on this 275 labeled frame dataset with a ResNet50-based neural network for 250,000 iterations. After evaluating the network, the train error was calculated at 1.24 pixels, and the test error was measured at 1.25 pixels. Once the model was successfully trained, all behavioral videos of the five mice were analyzed. To estimate pupil area, a python library (scikit-image)^172^ was used to fit an ellipse on the estimations of the eight markers generated by DeepLabCut. This ellipse model was used to predict the values of the vertices and co-vertices on the ellipse, permitting the calculation of the major and minor axes. Thus, these measurements were used to approximate the area of the pupil for each frame. To detect blinking behavior, the maximum pupil size in non-blinking conditions was calculated and used as a threshold to identify abnormally high pupil predictions. As such, pupil areas greater than 450 pixels were removed via interpolation. The pupil area data were saved as a csv file for importing into MATLAB.

### Fiber Photometry Signal Processing

Custom-written MATLAB scripts were used for post processing of the fluorescent signals. Photobleaching was corrected in both the isosbestic and GCaMP signals using detrended lines of best fit and subtracting the line from all values. After, the isosbestic and GCaMP median absolute deviation of z-score (ZMAD) signals were calculated before subtracting the GCaMP signal from the isosbestic to remove calcium-independent artifacts.

### Alignment of Task-Related Components

A custom-written MATLAB script imported the lick-related and overall responding data, the processed ZMAD GCaMP signal, and the processed pupil csv for alignment. The overall responding data (containing trial elements such as time of PT window start which is the TTL flag within Synapse) was converted from UNIX timecode into seconds to match the Synapse (containing the pupil video and processed GCaMP signal) time. Pupil data was resampled to align with GCaMP signal using rational fraction approximation. Furthermore, the length of each trial, as determined by the overall responding data, was recorded and used to capture the GCaMP and pupil data within each trial window. Licks were identified by setting upper and lower thresholds, and detecting when either was crossed. At this point, all parameters (e.g., GCaMP, pupil, and licking activity) were captured and aligned within the time window of each trial. These parameters could now be segmented by trial type (e.g., Go vs. NoGo texture) and by trial outcome (e.g., Hit, Miss, FA, CR) for more advanced analysis.

### Calcium Analysis

auROC is an analysis commonly applied to calcium imaging data to characterize the stereotypy of neuronal responses.^121,122^ Briefly, a baseline window is set within a non-task-related component of the overall calcium signal and compared to the rest of the signal via a sliding window. The maximal value of each signal is captured at each behavioral time point and averaged across the cohort. auROC values equal to 0.50 indicate no differences between the baseline signal and the task-related signal.

To assess longitudinal changes in calcium activity, two windows were established: a Control Window located from trial start to the sound cue indicating trial start, and a Target Window located 2s before and after PT start (overall = 4s). For each trial, all calcium peaks were measured, and onl^y^ peaks ≥ 90th percentile with a minimum peak prominence of 2 were selected. Finally, the maximum values within the Control and Target Windows (if present) were captured and averaged for every session.^173^

### Statistical Analyses

All data are reported as mean ± SEM unless otherwise noted. Statistical analyses were performed in GraphPad Prism (USA). All data were tested using the Shapiro-Wilk normality test. The means of different data distributions were analyzed and compared using two-tailed Student’s t-test (**Figures S1O, S1R, 2E FA Rate, 2E Bias, 3F, 4E Hit Rate, 4E Sensitivity, 4E Bias, 4G FA Rate, 4G Bias, 4K Learning/Expert Hit Rate, 4K Learning/Expert FA Rate, 4K Learning/Expert Sensitivity, 4K Learning/Expert Bias, 4L, S5C Hit Rate, S5C FA Rate, S5C Sensitivity, S5C Bias**), Wilcoxon signed rank test (**Figures S1N, S1Q, 2E Hit Rate, 2E Sensitivity, 4E FA Rate, 4G Hit Rate, 4G Sensitivity**), ordinary one-way ANOVA with Tukey’s multiple comparison correction (**Figures S1G, S1L**), Kruksal-Wallis with Dunn’s multiple comparison correction (**Figures 1E, 1F, 1I, 1L, S1H, S1I, S1J, S1K, 4I**), linear regression (**Figure S1E**), Repeated measures one-way ANOVA with Tukey’s multiple comparison correction (**Figures 3B, 3C**), Repeated measures mixed-effects analysis with Tukey’s multiple comparison correction (**Figures S4D Top, S4D Bottom**), Repeated measures two-way ANOVA with Sidak’s multiple comparison correction (**Figures 3K, 3L**), Repeated measures mixed-effects analysis with Sidak’s multiple comparison correction (**Figure 3Q**). For all statistical tests, *p < 0.05, **p < 0.01, ***p < 0.001, and ****p < 0.0001.

## Supplemental Data

**Figure S1.**
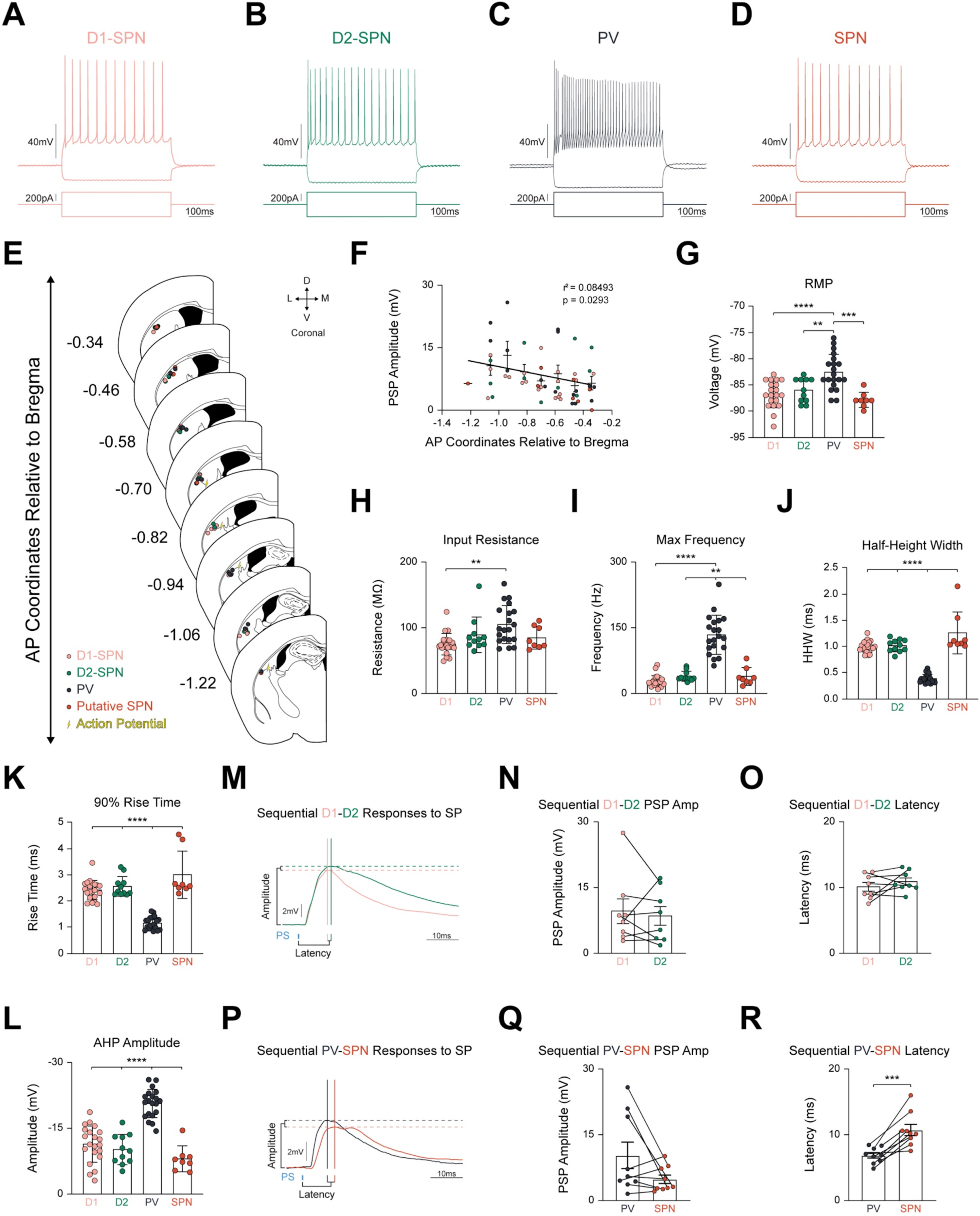
Recorded Cell Location, Intrinsic Electrophysiological Parameters, and Sequentially Patched PSP Amplitude and Latency. Related to Figure 1. **(A-D)** Representative responses of **(A)** D1-SPN (salmon), **(B)** D2-SPN (green), **(C)** PV interneuron (dark gray), and **(D)** unidentified SPN (orange) to hyperpolarizing and depolarizing current injections. Vertical voltage scale = 40mV. Horizontal current scale = 100ms. Vertical current scale = 50pA. **(E)** Recording location schematic of all recorded cells including cells excluded due to action potentials (lightning bolt). Note = all recordings took place within posterior DLS as it is the only striatal region innervated by POm axonal projections [S1, S2]. **(F)** Small but significant correlation (r^2^ = 0.08493; p = 0.0293) between PSP amplitude and increasing distance from the injection site (AP = -2.05) excluding cells with action potentials. **(G-L)** Parameters are consistent with literature concerning both SPNs and PV interneurons: **(G)** Mean resting membrane potential (RMP; D1-SPN: -86.80±0.58mV; D2-SPN: -86.00±0.69mV; PV: - 82.55±0.76mV; SPN: -87.88±0.52mV) (F_(4,56)_ = 11.24, p < 0.0001, D1 vs. PV p < 0.0001, D2 vs. PV p = 0.0079, SPN vs. PV p = 0.0001), **(H)** Mean input resistance (Rin; D1: 74.97±3.60MΩ, D2: 89.25±8.25MΩ, PV: 105.00±6.51MΩ, SPN: 84.58±6.16MΩ) (F_(4,56)_ = 14.27, p = 0.0026, D1 vs. PV p = 0.0011, **(I)** Mean maximum frequency (D1: 26.34±2.95Hz, D2: 38.38±3.46Hz, PV: 134.00±10.07Hz, SPN: 38.00±7.25Hz) (F_(4,56)_ = 42.64, p < 0.0001, D1 vs. PV p < 0.0001, D2 vs. PV p = 0.0032, SPN vs. PV p = 0.0027), **(J)** Mean half-height width (HHW; D1: 0.99±0.02ms, D2: 1.01±0.04ms, PV: 0.38±0.02ms, SPN: 1.26±0.14ms) (F_(4,56)_ = 42.06, p < 0.0001, D1 vs. PV p < 0.0001, D2 vs. PV p < 0.0001, SPN vs. PV p < 0.0001), **(K)** Mean 90% rise time (D1: 2.42±0.08ms, D2: 2.56±0.12ms, PV: 1.14±0.05ms, SPN: 3.00±0.32ms) (F_(4,56)_ = 40.52, p < 0.0001, D1 vs. PV p < 0.0001, D2 vs. PV p < 0.0001, SPN vs. PV p < 0.0001), and **(L)** Mean afterhyperpolarization (AHP) amplitude (D1: -11.02±0.91mV, D2: -9.79±1.05mV, PV: -20.39±0.73mV, SPN: -7.64±1.05mV) (F_(4,56)_ = 38.28, p < 0.0001, D1 vs. PV p < 0.0001, D2 vs. PV p < 0.0001, SPN vs. PV p < 0.0001). **(M-O)** Identified and unidentified neurons were patched sequentially and tested using the same protocols (SP, PPR, and Train) to control for injection site variability. **(M)** Representative PSP responses of a sequentially patched D1- and D2-SPN pairing (n = 8 D1-D2-SPN sequential recordings from 5 mice). Time scale = 10ms. Voltage scale = 2mV. **(N)** No significant differences in PSP amplitude between sequentially patched D1-SPN and D2-SPN pairs. **(O)** No significant differences in latency between sequentially patched D1-SPN and D2-SPN pairs. **(P)** Same as in **M** for recording acquired from sequentially patched PV and SPN pairings (n = 9 PV-SPN sequential recordings from 7 mice). Time scale = 10ms. Voltage scale = 2mV. **(Q)** No significant difference in PSP amplitude between sequentially patched PV and SPN pairs. **(R)** Significant differences in latency between sequentially patched PV and SPN pairs (6.85±0.42 for PV vs. 10.75±0.87 for SPN, p = 0.0006, n = 9 pairs). Data are mean ± SEM. ** p < 0.01, *** p < 0.001, **** p < 0.0001.

**Figure S2.**
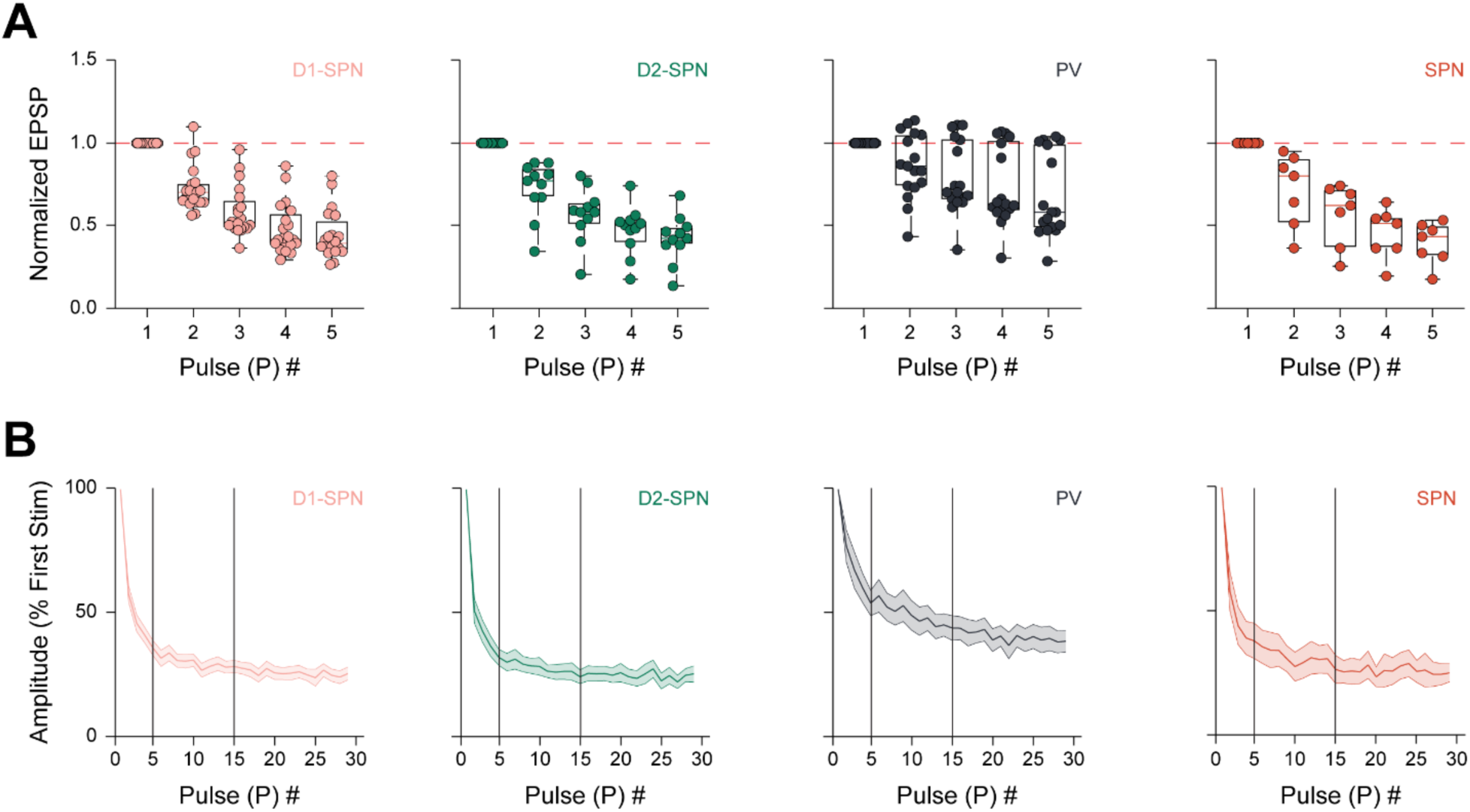
Representative Responses of Identified Striatal Cells to PPR and Train Stimulation. Related to Figure 1. **(A)** Box-and-whisker plot of the normalization of the second PSP to the first PSP of all recorded D1-SPNs (salmon), D2-SPNs (green), PV interneurons (dark gray), and unidentified SPNs (orange). **(B)** Summary plots of all PSP amplitudes normalized to the first pulse for all recorded D1-SPNs, D2-SPNs, PV interneurons, and unidentified SPNs. Vertical black lines indicate the averaging within from pulse 5 to pulse 15. Data are mean ± SEM.

**Figure S3.**
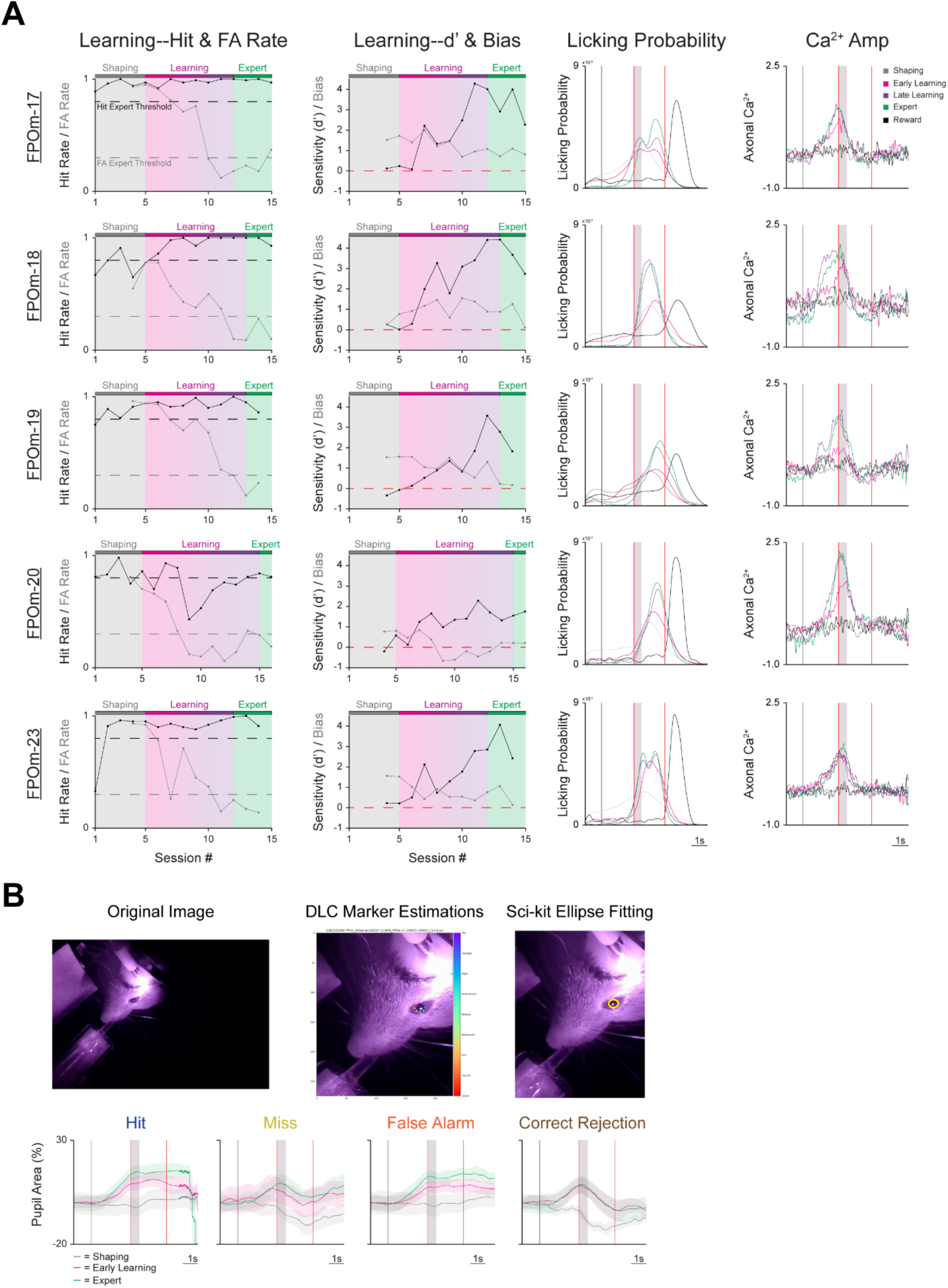
Individual Longitudinal Learning-Related Changes in Behavioral Parameters, Licking Activity, and Calcium Activity, and Methodology of Measuring Pupil Dynamics. Related to Figure 2. **(A)** Longitudinal learning-related changes as mice progress through each time point (Shaping, Early Learning, Late Learning, and Expert). The Shaping time point permitted mice to learn to lick the spout for a water reward during the first three sessions without the presence of the textures. During the last two Shaping sessions, the Go and NoGo textures were presented simultaneously with a Go texture probability of 0.90 for the first session and 0.75 for the second session. For the first Learning session, and all subsequent sessions, the Go texture probability was set to 0.50. Mice were considered Learning at this time point. The Early Learning time point was considered the first two sessions after Shaping, while the Late Learning was considered the last two sessions before achieving Expert status. Mice were considered Expert when they had a Hit Rate ≥ 0.80 and a FA Rate ≤ 0.30 for two consecutive sessions. After discrimination training was completed, mice were subjected to a Reward session. During the Reward session, the lick thresholds were unobtainable, the textures were oriented so they could not be contacted by the whiskers, and water was automatically delivered at the end of the PT window (second vertical red line). *Left*: Hit Rate (black) and FA Rate (gray) across learning. *Left Middle*: Sensitivity (d’; black) and Bias (gray) across learning. Red dashed line indicates 0. Note that a strict d’ threshold was not used due to artificially increased d’ values as Hit Rate and/or FA Rate approached their extremes (0 or 1) as in FPOm-18 sessions 8 to 15. *Right Middle*: Probability density function for licking activity at each time point. *Right*: Average axonal ZMAD calcium activity at each time point. Scale bar = 1s. **(B)** To measure pupil dynamics during the Go/NoGo discrimination task, orofacial video was synchronously recorded with calcium activity. This video was cropped in DeepLabCut [S3, S4], and nine markers were manually placed within the video: eight circumscribing the pupil (top, top right, right, bottom right, bottom, bottom left, left, and top left) and one on the spout. Once the model was trained, an ellipse was fitted through the eight pupil markers in sci-kit [S5]. Pupil values were converted from pixels to percentage, and the baseline was normalized to 0. **(B)** Notably, pupil-related dynamics (e.g. Shaping, Early Learning and Expert) during the trial outcomes (e.g. Hit, Miss, False Alarm, and Correct Rejection) are similar to previously observed dynamics [S6] despite using a deep learning-based methodology [S7]. Data are mean ± SEM. Scale bar = 1s.

**Figure S4.**
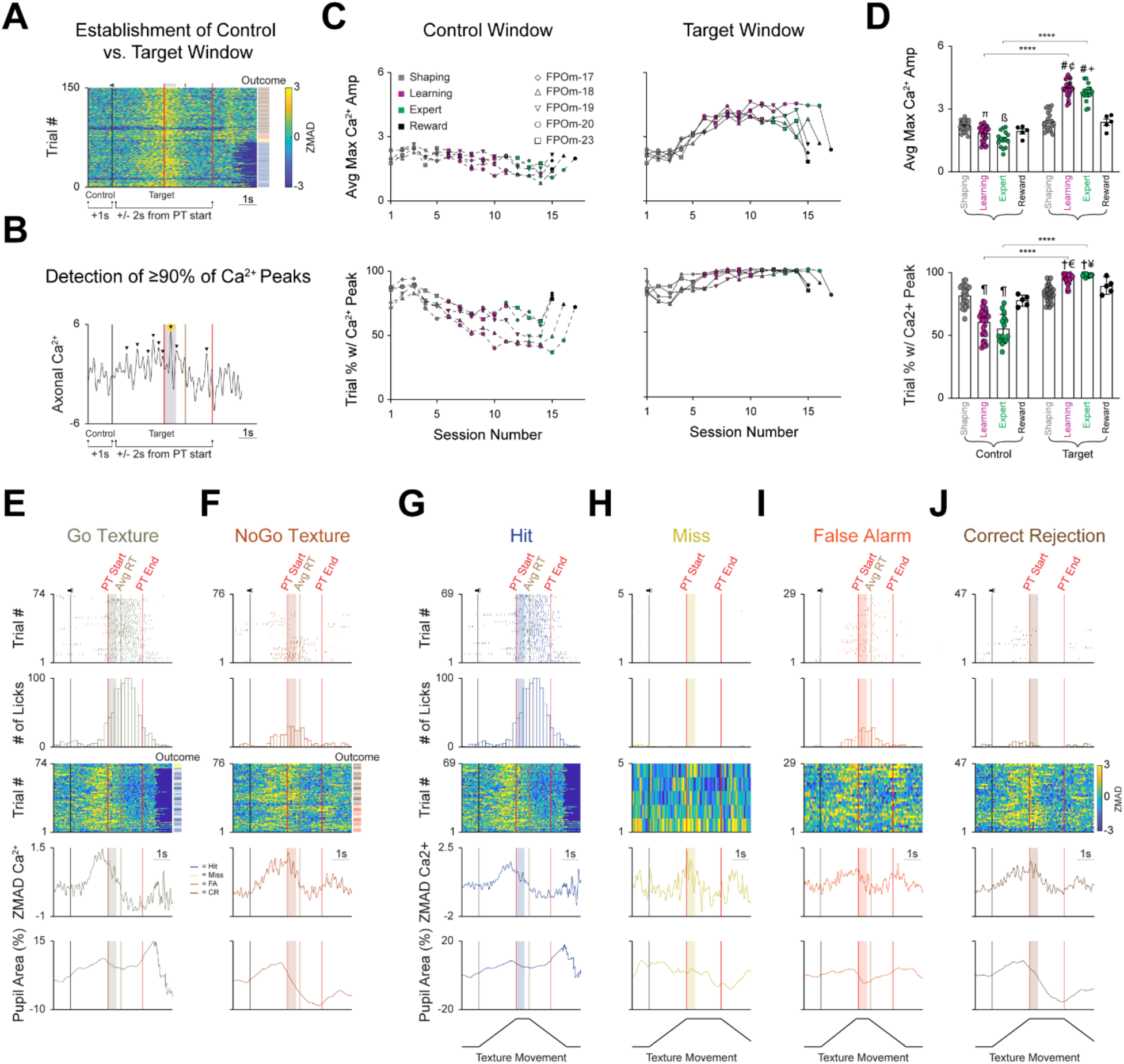
Establishment of Control and Target Windows, and Representative Example of All Three Activity Parameters Segmented by Trial Type and Outcome. Related to Figure 3. **(A)** Establishment of the Control (1 second following trial start, encompassing the pre-trial intertrial interval) and Target (2 seconds before and after PT window start) windows. Task-related events are directly above the heatmap. Trial outcomes are color coded (blue = Hit, yellow = Miss, orange = False Alarm (FA), brown = Correct Rejection (CR). **(B)** Representative axonal calcium activity during a single trial (outcome = hit). Black arrowheads above positive deflections indicate a calcium peak that was greater than or equal to the 90th percentile of all calcium peaks. The yellow circle encompassing the largest calcium peak was selected as the maximal calcium amplitude. **(C)** Longitudinal representation of the average maximum calcium amplitude (*Top*) and the average trial % with a detectable calcium peak (*Bottom*) within the Control (*Left*) and Target (*Right*) windows. **(D)** Average maximum calcium amplitude (*Top*, ᴨ p = 0.0005, Control Shaping vs. Control Learning; ß p < 0.0001, Control Shaping vs. Control Expert; # p < 0.0001, Target Shaping vs. Target Learning/Expert; ¢ p = 0.0073, Target Learning vs. Target Reward; + p = 0.0091 Target Expert vs. Target Reward; Mixed-Effects Analysis with Tukey’s correction for multiple comparisons). Average trial % with a detectable calcium peak (*Bottom*, ¶ p < 0.0001 Control Shaping vs. Control Learning/Expert; † p < 0.0001, Target Shaping vs. Target Learning/Expert; € p = 0.0073, Target Learning vs. Target Reward; ¥ p = 0.0091, Target Expert vs. Target Reward; Mixed-Effects Analysis with Tukey’s correction for multiple comparisons). **(E-F)** Three activity parameters (licking, axonal calcium, and pupil) from a representative session segmented by trial type: **(E)** Go texture or **(F)** NoGo texture presentation. *Top*: Licking activity. Licks are denoted as colored tick marks. The vertical black line represents a sound cue indicating trial start as the presented texture moves towards the whisker field. The vertical red lines denote the start (texture arrival at its endpoint in the whisker field) and end of the PT window. The vertical brown line denotes average RT. The colored boxes denote a 500ms grace period wherein the mouse can lick freely without triggering any outcomes. *Top Middle*: Lick histogram. *Middle*: Heatmap sorted by trial outcome (*Right* of heatmap) highlighting axonal calcium activity for each trial. Trial outcome is color coded (Blue = Hit, yellow = Miss, Orange = FA, Brown = CR). *Bottom Middle*: Average axonal calcium activity. *Bottom*: Average pupil area (as a normalized percentage). **(G-J)** The representative session from E-F was further segmented by trial outcome: **(G)** Hit, **(H)** Miss, **(I)** FA, and **(J)** CR. Underneath the *Bottom* panel, texture movement that is dependent on trial outcome is illustrated. Data are mean ± SEM. **** p < 0.0001.

**Figure S5.**
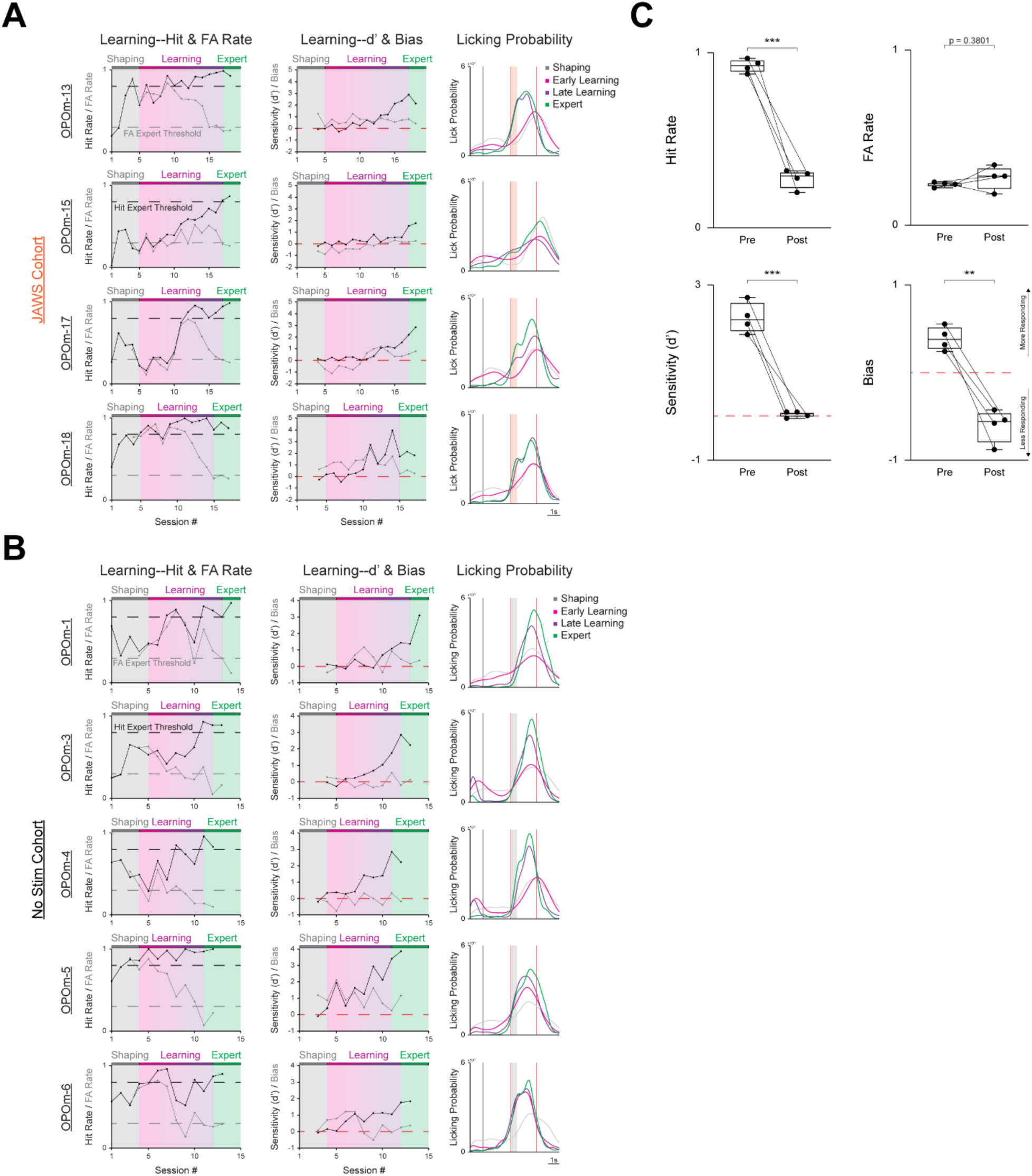
Individual Longitudinal Learning-Related Changes in Behavioral Parameters and Licking Activity for the JAWS and No Stim cohorts, and Whisker Trim. Related to Figure 4. **(A-B)** Longitudinal learning-related changes as mice from the **(A)** JAWS and the No Stim **(B)** cohorts progress through each behavioral time point (Shaping, Early Learning, Late Learning, and Expert). The Shaping time point permitted mice to learn to lick the spout for a water reward during the first three sessions without the presence of the textures. During the last two Shaping sessions, the Go and NoGo textures were presented simultaneously with a Go texture probability of 0.90 for the first session and 0.75 for the second session. For the first Learning session, and all subsequent sessions, the Go texture probability was set to 0.50. Mice were considered Learning at this time point. The Early Learning time point was considered the first two sessions after Shaping, while the Late Learning was considered the last two sessions before achieving Expert status. Mice were considered Expert when they had a Hit Rate ≥ 0.80 and a FA Rate ≤ 0.30 for two consecutive sessions. *Left*: Hit Rate (black) and FA Rate (gray) across learning. *Middle*: Sensitivity (d’; black) and Bias (gray) across learning. Red dashed line indicates 0. Note that a strict d’ threshold was not used due to artificially increased d’ values as Hit Rate and/or FA Rate approached their extremes (0 or 1). *Right*: Probability density function for licking activity at each time point. Scale = 1s. **(C)** Following the attainment of expert status, a group of mice (n = 4) underwent a whisker trim session. In this session, mice performed 75 trials as normal with no optogenetic stimulation. After, the right whiskers contacting the textures were trimmed, and another 75 trials were performed. Behavioral performance, based on the four parameters (Hit Rate, FA Rate, Sensitivity, and Bias), was compared. Data are mean ± SEM. ** p < 0.01, *** p < 0.001.

## References

1. Petersen, C.C.H. (2019). Sensorimotor processing in the rodent barrel cortex. Nat Rev Neurosci, 20: 533–546.

2. El-Boustani, S., Sermet, B.S., Foustokos, G., Oram, T.B., Yizhar, O., Petersen, C.C.H. (2020). Anatomically and functionally distinct thalamocortical inputs to primary and secondary mouse whisker somatosensory cortices. Nat Commun, 11(3342).

3. Deschenes, M., Timofeeva, E., Lavallee, P. (2003). The relay of high-frequency sensory signals in the whisker-to-barreloid pathway. J Neurosci, 23: 6778–6787.

4. Bureau, I., von Saint Paul, F., Svoboda, K. (2006). Interdigitated paralemniscal and lemniscal pathways in the mouse barrel cortex. PLoS Biol, 4(12): e382.

5. Petersen, C.C.H. (2007). The functional organization of the barrel cortex. Neuron, 56: 339–355.

6. Diamond, M.E., von Heimendalh, M., Knutsen, P.M., Kleinfeld, D., Ahissar, E. (2008). ‘Where’ and ‘what’ in the whisker sensorimotor system. Nat Rev Neurosci, 9: 601–612.

7. Moore, J.D., Lindsay, N.M., Deschenes, M., Kleinfeld, D. (2015). Vibrissa self-motion and touch are reliably encoded along the same somatosensory pathway from brainstem through thalamus. PLoS Biol, 13: e1002253.

8. Deschenes, M., Urbain, N. (2016). Vibrissal afferents from trigeminus to cortices. In Prescott, TJ et al. (eds), Scholarpedia of Touch, Atlantis press.

9. Yu, C., Derdikman, D., Haidarliu, S., Ahissar, E. (2006). Parallel thalamic pathways for whisking and touch signals in the rat. PLoS Biol, 4(5): e124.

10. Mo, C., Petrof, I., Viaene, A.N., Sherman, S.M. (2017). Synaptic properties of the lemniscal and paralemniscal pathway to the mouse somatosensory thalamus. Proc Natl Acad Sci USA, 114(30): e6212–e6221.

11. Chiaia, N.L., Rhoades, R.W., Fish, S.E., Killackey, H.P. (1991a). Thalamic processing of vibrissal information in the rat: II. Morphological and functional properties of medial ventral posterior and posterior nucleus neurons. J Comp Neurol, 314: 217–236.

12. Diamond, M.E., Armstrong-James, M., Ebner, F.F. (1992). Somatic sensory responses in the rostral sector of the posterior group (POm) and in the ventral posterior medial nucleus (VPM) of the rat thalamus. J Comp Neurol, 318: 462–476.

13. Sosnik, R., Haidarliu, S., Ahissar, E. (2001). Temporal frequency of whisker movement: I. representations in brain stem and thalamus. J Neurophysiol, 86: 339–353.

14. Urbain, N., Salin, P.A., Libourel, P.A., Comte, J.C., Gentet, L.J., Petersen, C.C.H. (2015). Whisking-related changes in neuronal firing and membrane potential dynamics in the somatosensory thalamus of awake mice. Cell Reports, 13: 647–656.

15. Chiaia, N.L., Rhoades, R.W., Bennett-Clarke, C.A., Fish, S.E., Killackey, H.P. (1991b). Thalamic processing of vibrissal information in the rat: I. afferent input to the medial ventral posterior and posterior nuclei. J Comp Neurol, 314: 217–236.

16. Lavallee, P., Urbain, N., Dufresne, C., Bokor, H., Acsady, L., Deschenes, M. (2005). Feedforward inhibitory control of sensory information in higher-order thalamic nuclei. J Neurosci, 25(33): 7489–7498.

17. Masri, R., Bezdudnaya, T., Trageser, J.C., Keller, A. (2008). Encoding of stimulus frequency and sensor motion in the posterior medial thalamic nucleus. J Neurophysiol, 100: 681–689.

18. Petty, G.H., Kinnischtzke, A.K., Kate Hong, Y., Bruno, R.M. (2021). Effects of arousal and movement on secondary somatosensory and visual thalamus. eLife, 10: e67611.

19. Noseda, R., Kainz, V., Jakubowski, M., Gooley, J.J., Saper, C.B., Digre, K., Burstein, R. (2010). A neural mechanism for exacerbation of headache by light. Nat Neurosci, 13: 239–245.

20. Frangeul, L., Porrero, C., Garcia-Amado, M., Maimone, B., Maniglier, M., Clasca, F., Jabaudon, D. (2014). Specific activation of the paralemniscal pathway during nociception. Eur J Neurosci, 39: 1455–1464.

21. Osaki, H., Kanaya, M., Ueta, Y., Miyata, M. (2022). Distinct nociception processing in the dysgranular and barrel regions of the mouse somatosensory cortex. Nat Commun, 13(3622).

22. Gambino, F., Pages, S., Kehayas, V., Baptista, D., Tatti, R., Carleton, A., Holtmaat, A. (2014). Sensory-evoked LTP driven by dendritic plateau potentials in vivo. Nature, 515: 116–119.

23. Audette, N.J., Bernhard, S.M., Ray, A., Stewart, L.T., Barth, A.L. (2019). Rapid plasticity of higher-order thalamocortical inputs during sensory learning. Neuron, 103: 277–291.

24. Williams, L.E., Holtmaat, A. (2019). Higher-order thalamocortical inputs gate synaptic long-term potentiation via disinhibition. Neuron, 101: 91–102.

25. Zhang, W., Bruno, R.M. (2019). High-order thalamic inputs to primary somatosensory cortex are stronger and longer lasting than cortical inputs. eLife, 8: e44158.

26. La Terra, D., Bjerre, A.S., Rosier, M., Masuda, R., Ryan, T.J., Palmer, L.M. (2022). The role of higher-order thalamus during learning and correct performance in goal-directed behavior. eLife, 11: e77177.

27. Roger, M., Cadusseau, J. (1984). Afferent connections of the nucleus posterior thalami in the rat, with some evolutionary and functional considerations. J Hirnforsch, 25: 473–485.

28. Pierret, T., Lavallee, P., Deschenes, M. (2000). Parallel streams for the relay of vibrissal information through thalamic barreloids. J Neurosci, 20: 7455–7462.

29. Alloway, K.D., Hoffer, Z.S., Hoover, J.E. (2003). Quantitative comparisons of corticothalamic topography within the ventrobasal complex and the posterior nucleus of the rodent thalamus. Brain Res, 968: 54–68.

30. Alloway, K.D., Olson, M.L., Smith, J.B. (2008). Contralateral corticothalamic projections M1 whisker cortex: potential route for modulating hemispheric interactions. J Comp Neurol, 510(1): 100–116.

31. Groh, A., Bokor, H., Mease, R.A., Plattner, V.M., Hangya, B., Stroh, A., Deschenes, M., Acsady, L. (2014). Convergence of cortical and sensory driver inputs on single thalamocortical cells. Cereb Cortex, 24(12): 3167–3179.

32. Yamawaki, N., Shepherd, G.M.G. (2015). Synaptic circuit organization of motor corticothalamic neurons. J Neurosci, 35(5): 2293–2307.

33. Sumser, A., Mease, R.A., Sakmann, B., Groh, A. (2017). Organization and somatotopy of corticothalamic projections from L5b in mouse barrel cortex. Proc Natl Acad Sci USA, 114(33): 8853–8858.

34. Gharaei, S., Honnuraiah, S., Arabzadeh, E., Stuart, G.J. (2020). Superior colliculus modulates cortical coding of somatosensory information. Nat Commun, 11(1693).

35. Roger, M., Cadusseau, J. (1985). Afferents to the zona incerta in the rat: a combined retrograde and anterograde study. J Comp Neurol, 241: 480–492.

36. Power, B.D., Kolmac, C.I., Mitrofanis, J. (1999). Evidence for a large projection from the zona incerta to the dorsal thalamus. J Comp Neurol, 404: 554–565.

37. Bartho, P., Freund, T.F., Acasady, L. (2002). Selective GABAergic innervation of thalamic nuclei from zona incerta. Eur J Neurosci, 16: 999–1014.

38. Power, B.D., Mitrofanis, J. (2002). Ultrastructure of afferents from the zona incerta to the posterior and parafascicular thalamic nuclei of rats. J Comp Neurol, 451: 33–44.

39. Trageser, J.C., Keller, A. (2004). Reducing the uncertainty: gating of peripheral inputs by zona incerta. J Neurosci, 24: 8911–8915.

40. Bartho, P., Slezia, A., Varga, V., Bokor, H., Pinault, D., Buzaski, G., Acsady, L. (2007). Cortical control of zona incerta. Eur J Neurosci, 27(7): 1670–1681.

41. Giber, K., Slezia, A., Bokor, H., Bodor, A.L., Ludanyi, A., Katona, I., Acsady, L. (2008). Heterogeneous output pathways link the anterior pretectal nucleus with the zona incerta and the thalamus in the rat. J Comp Neurol, 506(1): 122–140.

42. Watson, G.D.R., Smith, J.B., Alloway, K.D. (2015). The zona incerta regulates communication between the superior colliculus and the posteromedial thalamus: implications for thalamic interactions with the dorsolateral striatum. J Neurosci, 35: 9463–9476.

43. Masri, R., Trageser, J.C., Bezdudnaya, T., Li, Y., Keller, A. (2006). Cholinergic regulation of the posterior medial thalamic nucleus. J Neurophysiol, 96(5): 2256–2273.

44. Trageser, J.C., Burke, K.A., Masri, R., Li, Y., Sellers, L., Keller, A. (2006). State-dependent gating of sensory inputs by zona incerta. J Neurophysiol, 96: 1456–1463.

45. Park, A., Li, Y., Masri, R., Keller, A. (2017). Presynaptic and extrasynaptic regulation of posterior nucleus of thalamus. J Neurophysiol, 118(1): 507–519.

46. Huerto-Ocampo, I., Hacioglu-Bay, H., Dautan, D., Mena-Segovia, J. (2019). Distribution of midbrain cholinergic axons in the thalamus. eNeuro, 7(1): eneuro.0454-19.2019.

47. Meyer, H.S., Wimmer, V.C., Hemberger, M., Bruno, R.M., de Kock, C.P.J., Frick, A., Sakmann, B., Helmstaedter, M. (2010). Cell type-specific thalamic innervation in a column of rat vibrissal cortex. Cereb Cortex, 20: 2287–2303.

48. Wimmer, V.C., Bruno, R.M., de Kock, C.P.J., Kuner, T., Sakmann, B. (2010). Dimensions of a projection column and architecture of VPM and POm axons in rat vibrissal cortex. Cereb Cortex, 20(10): 2265–2276.

49. Ohno, S., Kuramoto, E., Furuta, T., Hioki, H., Tanaka, Y.R., Fujiyama, F., Sonomura, T., Uemura, M., Sugiyama, K., Kaneko, T. (2012). Morphological analysis of thalamocortical axon fibers of rat posterior thalamic nuclei: a single neuron tracing study with viral vectors. Cereb Cortex, 22(12): 2840–2857.

50. Audette, N.J., Urban-Ciecko, J., Matsushita, M., Barth, A.L. (2018). POm thalamocortical input drives layer-specific microcircuits in somatosensory cortex. Cereb Cortex, 28: 1312–1328.

51. Qi, J., Ye, C., Naskar, S., Inacio, A.R., Lee, S. (2022). Posteromedial thalamic nucleus activity significantly contributes to perceptual discrimination. PLoS Biol, 20(11): e3001896.

52. Spreafico, R., Barbaresi, P., Weinberg, R.J., Rustioni, A. (1987). SII-projecting neurons in the rat thalamus: a single- and double-retrograde-tracing study. Somatosens Res, 4: 359–357.

53. Deschenes, M., Bourassa, J., Parent, A. (1995). Two different types of thalamic fibers innervate the rat striatum. Brain Res, 701: 288–292.

54. Viaene, A.N., Petrof, I., Sherman, S.M. (2011). Properties of the thalamic projection from the posterior medial nucleus to primary and secondary somatosensory cortices in the mouse. Proc Natl Acad Sci USA, 108: 18156–18161.

55. Smith, J.B., Mowery, T.M., Alloway, K.D. (2012). Thalamic POm projections to the dorsolateral striatum of rats: potential pathway for mediating stimulus-response associations for sensorimotor habits. J Neurophysiol, 108: 160–174.

56. Alloway, K.D., Smith, J.B., Mowery, T.M., Watson, G.D.R. (2017). Sensory processing in the dorsolateral striatum: the contribution of thalamostriatal pathways. Front Syst Neurosci, 11: 53.

57. Li, Y., Lopez-Huerta, V.G., Adiconis, X., Levandowski, K., Choi, S., Simmons, S.K., Arias-Garcia, M.A., Guo, B., Yao, A.Y., Blosser, T.R., et al. (2020). Distinct subnetworks of the thalamic reticular nucleus. Nature, 583’ 819-824.

58. O’Reilly, C., Iavarone, E., Yi, J., Hill, S.L. (2021). Rodent somatosensory thalamocortical circuitry: neurons, synapses, and connectivity. Neurosci Biobehav Rev, 126: 213–235.

59. Yonk, A. J., Margolis, D. J. (2019). Traces of learning in thalamocortical circuits. Neuron, 103(2): 175–176.

60. Gerfen, C.R., Wilson, C.J. (1996). “The basal ganglia” in Handbook of Chemical Neuroanatomy, 12^th^ Ed, eds Swanson, L.W., Bjorklund, A., Hokfelt, T (Amsterdam: Elsevier Science), 371-468.

61. Gerfen, C.R., Surmeier, D.J. (2011). Modulation of striatal projection systems by dopamine. Ann Rev Neurosci, 34: 441–466.

62. Tritsch, N.X., Sabatini, B.L. (2012). Dopaminergic modulation of synaptic transmission in cortex and striatum. Neuron, 76(1): 33–50.

63. Gittis, A.H., Nelson, A.B., Thwin, M.T., Palop, J.J., Kreitzer, A.C. (2010). Distinct roles of GABAergic interneurons in the regulation of striatal output pathways. J Neurosci, 30: 2223–2234.

64. Tepper, J.M., Tecuapetla, F., Koos, T., Ibanez-Sandoval, O. (2010). Heterogeneity and diversity of striatal GABAergic interneurons. Front Neuroanat, 4: 150.

65. Lee, K., Holley, S.M., Shobe, J.L., Chong, N.C., Cepeda, C., Levine, M.S., Masmanidis, S.C. (2017). Parvalbumin interneurons modulate striatal output and enhance performance during associative learning. Neuron, 93(6): 1451–1463.e4.

66. Owen, S.F., Berke, J.D., Kreizter, A.C. (2018). Fast-spiking interneurons supply feedforward control of bursting, calcium, and plasticity for efficient learning. Cell, 172: 683–695.

67. Tepper, J.M., Koos, T., Ibanez-Sandoval, O., Tecuapetla, F., Faust, T.W., Assous, M. (2018). Heterogeneity and diversity of striatal GABAergic interneurons: update 2018. Front Neuroanat, 12: 91.

68. Pan, W., Mao, T., Dudman, J.T. (2010). Inputs to the dorsal striatum of the mouse reflect the parallel architecture of the forebrain. Front Neuroanat, 4: fnana.2010.00147.

69. Wall, N.R., De La Parra, M., Callaway, E.M., Kreitzer, A.C. (2013). Differential innervation of direct- and indirect-pathway striatal projection neurons. Neuron, 79(2): 347–360.

70. Huerto-Ocampo, I., Mena-Segovia, J., Bolam, J.P. (2014). Convergence of cortical and thalamic input to direct and indirect pathway medium spiny neurons in the striatum. Brain Struct Funct, 219(5): 1787–1800.

71. Oh, S.W., Harris, J.A., Ng, L., Winslow, B., Cain, N., Mihalas, S., Wang, Q., Lau, C., Kuan, L., Henry, A.M., et al. (2014). A mesoscale connectome of the mouse brain. Nature, 508: 207–214.

72. Guo, Q., Wang, D., He, X., Feng, Q., Lin, R., Xu, F., Fu, L., Luo, M. (2015). Whole-brain mapping of inputs to projection neurons and cholinergic interneurons in the dorsal striatum. PLoS One, 10(4): e0123381.

73. Hunnicutt, B.J., Jongbloets, B.C., Birdsong, W.T., Gertz, K.J., Zhong, H., Mao, T. (2016). A comprehensive excitatory input map of the striatum reveals novel functional organization. *eLife*, e19103.

74. Hintiryan, H., Foster, N.N., Bowman, I., Bay, M., Song, M.Y., Gou, L., Yamashita, S., Bienkowski, M.S., Zingg, B., Zhu, M., et al. (2016). The mouse cortico-striatal projectome. Nat Neurosci, 19(8): 1100–1114.

75. Smith, J.B., Klug, J.R., Ross, D.L., Howard, C.D., Hollon, N.G., Ko, V.I., Hoffman, H., Callaway, E.M., Gerfen, C.R., Jin, X. (2016). Genetic-based dissection unveils the inputs and outputs of striatal patch and matrix compartments. Neuron, 91: 1069–1084.

76. Hooks, B.M., Papale, A.E., Paletzki, R.F., Feroze, M.W., Eastwood, B.S., Couey, J.J., Winnubst, J., Chandrashekar, J., Gerfen, C.R. (2018). Topographic precision in sensory and motor corticostriatal projections varies across cell type and cortical area. Nat Commun, 9, 3549.

77. Alexander, G.E., DeLong, M.R., Strick, P.L. (1986). Parallel organization of functionally segregated circuits linking basal ganglia and cortex. Ann Rev Neurosci, 9: 357–381.

78. Mao, T., Kusefoglu, D., Hooks, B.M., Huber, D., Petreanu, L., Svoboda, K. (2011). Long-range neuronal circuits underlying the interaction between sensory and motor cortex. Neuron, 72: 111–123.

79. Chen, J.L., Carta, S., Soldado-Magraner, J., Schneider, B.L., Helmchen, F. (2013). Behaviour-dependent recruitment of long-range projection neurons in somatosensory cortex. Nature, 499: 336–340.

80. Kwon, S.E., Yang, H., Minamisawa, G., O’Connor, D.H. (2016). Sensory and decision-related activity propagate in a cortical feedback loop during touch perception. Nat Neurosci, 19: 1243–1249.

81. Alloway, K.D., Mutic, J.J., Hoffer, Z.S., Hoover, J.E. (2000). Overlapping corticostriatal projections from the rodent vibrissal representations in the primary and secondary somatosensory cortex. J Comp Neurol, 428: 51–67.

82. Hoffer, Z.S., Alloway, K.D. (2001). Organization of corticostriatal projections from the vibrissal representations in the primary motor and somatosensory cortical areas of rodents. J Comp Neurol, 439: 87–103.

83. Ramanathan, S., Hanley, J.J., Deniau, J.M., Bolam, J.P. (2002). Synaptic convergence of motor and somatosensory cortical afferents onto GABAergic interneurons in the rat striatum. J Neurosci, 22: 8158–8169.

84. Charpier, S., Pidoux, M., Mahon, S. (2020). Converging sensory and motor cortical inputs onto the same striatal neurons: an *in vivo* intracellular investigation. PLoS One, 15(2): e0228260.

85. Smith, J.B., Chakrabati, S., Mowery, T.M., Alloway, K.D. (2021). Convergence of forepaw somatosensory and motor cortical projections in the striatum, claustrum, thalamus, and pontine nuclei of cats. Brain Struct Funct, 227: 361–379.

86. Sanabria, B.D., Baskar, S.S., Yonk, A.J., Linares-Garcia, I., Abraira, V.E., Lee, C.R., Margolis, D.J. (2024). Cell-Type Specific Connectivity of Whisker-Related Sensory and Motor Cortical Input to Dorsal Striatum. eNeuro, 11(1): eneuro.0503-23.2023.

87. Ding, J., Peterson, J.D., Surmeier, D.J. (2008). Corticostriatal and thalamostriatal synapses have distinctive properties. J Neurosci, 28(25): 6483–6492.

88. Doig, N.M., Magill, P.J., Apicella, P., Bolam, J.P., Sharott, A. (2014). Cortical and thalamic excitation mediate the multiphasic responses of striatal cholinergic interneurons to motivationally salient stimuli. J Neurosci, 34: 3101–3117.

89. Lee, C.R., Yonk, A.J., Wiskerke, J., Paradiso, K.G., Tepper, J.M., Margolis, D.J. (2019). Opposing influence of sensory and motor cortical input on striatal circuitry and choice behavior. Curr Biol, 29: 1313–1323.e5.

90. Johansson, Y., Silberberg, G. (2020). The functional organization of cortical and thalamic inputs onto five types of striatal neurons is determined by source and target cell identities. Cell Reports, 30(4): 1178–1194.e3.

91. Sun, Z., Schneider, A., Alyahyay, M., Karvat, G., Diester, I. (2021). Effects of optogenetic stimulation on primary somatosensory cortex and its projections to striatum on vibrotactile perception in freely moving rats. eNeuro, 8(2): eneuro.0453-20.2021.

92. Zareian, B., Lam, A., Zagha, E. (2023). Dorsolateral striatum is a bottleneck for responding to task-relevant stimuli in a learned whisker detection task in mice. J Neurosci, 43(12): 2126–2139.

93. Smith, Y., Parent, A. (1986). Differential connections of the caudate nucleus and putamen in the squirrel monkey (saimiri sciureus). Neuroscience, 18: 347–371.

94. Berendse, H.W., Groenewegen, H.J. (1990). Organization of the thalamostriatal projections in the rat, with special emphasis on the ventral striatum. J Comp Neurol, 299: 187–228.

95. Lappar, S.R., Bolam, J.P. (1992). Input from the frontal cortex and the parafascicular nucleus to cholinergic interneurons in the dorsal striatum of the rat. Neuroscience, 51(3): 533–545.

96. Matsumoto, N., Minamimoto, T., Graybiel, A.M., Kimura, A. (2001). Neurons in the thalamic CM-Pf complex supply striatal neurons with information about behaviorally significant sensory events. J Neurophysiol, 85: 960–976.

97. Ellender, T.J., Harwood, J., Kosillo, P., Capogna, M., Bolam, J.P. (2013). Heterogeneous properties of lateral and parafascicular thalamic synapses in the striatum. J Physiol, 591: 257–272.

98. Parker, P.R.L., Lalive, A.L., Kreitzer, A.C. (2016). Pathway-specific remodeling of thalamostriatal synapses in Parkinsonian mice. Neuron, 89(4): 734–740.

99. Mandelbaum, G., Taranda, J., Haynes, T.M., Hochbaum, D.R., Huang, K.W., Hyun, M., Venkataraju, K.U., Straub, C., Wang, W., Robertson, K., et al. (2019). Distinct cortical-thalamic-striatal circuits through the parafascicular nucleus. Neuron, 102(3): 636–652.e7.

100. Tanimura, A., Du, Y., Kondapalli, J., Wokosin, D.L., Surmeier, D.J. (2019). Cholinergic interneurons amplify thalamostriatal excitation of striatal indirect pathway neurons in Parkinson’s disease models. Neuron, 101(3): 444–458.e6.

101. Fallon, I.P., Hughes, R.N., Ulloa Severino, F.P., Kim, N., Lawry, C.M., Watson, G.D.R., Roshchina, M., Yin, H.H. (2023). The role of the parafascicular thalamic nucleus in action initiation and steering. Curr Biol, 33: 1–11.

102. Brown, H.D., Baker, P.M., Ragozzino, M.E. (2010). The parafascicular thalamic nucleus concomitantly influences behavioural flexibility and dorsomedial striatal acetylcholine output in rats. J Neurosci, 30: 14390–14398.

103. Bradfield, L.A., Bertran-Gonzalez, J., Chieng, B., Balleine, B.W. (2013). The thalamostriatal pathway and cholinergic control of goal-directed action: interlacing new with existing learning in the striatum. Neuron, 79: 153–166.

104. Diaz-Hernandez, E., Contreras-Lopez, R., Sanchez-Fuentes, A., Rodriguez-Sibrian, L., Ramirez-Jarquin, J.O., Tecuapetla, F. (2018). The thalamostriatal projections contribute to the initiation and execution of a sequence of movements. Neuron, 100: 739–752.

105. Alloway, K.D., Smith, J.B., Watson, G.D.R. (2014). Thalamostriatal projections from the medial posterior and parafascicular nuclei have distinct topographic and physiologic properties. J Neurophysiol, 111: 36–50.

106. Kawaguchi, Y. (1993). Physiological, morphological, and histochemical characterization of three classes of interneurons in rat neostriatum. J Neurosci, 13(11): 4908–4923.

107. Abbott, L.F., Varela, J.A., Sen, K., Nelson, S.B. (1997). Synaptic depression and cortical gain control. Science, 275: 220–224.

108. Zucker, R.S., Regehr, W.G. (2002). Short-term synaptic plasticity. Ann Rev Physiol, 64: 355–405.

109. Assous, M., Tepper, J.M. (2019). Cortical and thalamic inputs exert cell-type specific feedforward inhibition on striatal GABAergic interneurons. J Neurosci Res, 97(12): 1491–1502.

110. Glasgow, S.D., McPhedrain, R., Madranges, J.F., Kennedy, T.E., Ruthazer, E.S. (2019). Approaches and limitations in the investigation of synaptic transmission and plasticity. Front Synapt Neurosci, 11: 20.

111. Landisman, C.E., Connors, B.W. (2007). VPM and POm nuclei of the rat somatosensory thalamus: intrinsic neuronal properties and corticothalamic feedback. Cereb Cortex, 17(12): 2853–2865.

112. Kawai, R., Markman, T., Poddar, R., Ko, R., Fantana, A.L., Dhawale, A.K., Kampff, A.R., Olveczky, B.P. (2015). Motor cortex is required for learning, but not for executing a motor skill. Neuron, 86: 800–812.

113. Rothwell, P.E., Hayton, S.J., Sun, G.L., Fuccillo, M.V., Lim, B.K., Malenka, R.C. (2015). Input- and output-specific regulation of serial order performance by corticostriatal circuits. Neuron, 88: 345–356.

114. Kupferschmidt, D.A., Juczewski, K., Cui, G., Johnson, K.A., Lovinger, D.M. (2017). Parallel, but dissociable, processing in discrete corticostriatal inputs encodes skill learning. Neuron, 96: 476–489.e5.

115. Kahneman, D., Beatty, J. (1966). Pupil diameter and load on memory. Science, 154, 1583–1585.

116. Murphy, P.R., Vandekerckhove, J., Nieuwenhuis, S. (2014). Pupil-linked arousal determines variability in perceptual decision making. PLoS Comput Biol, 10: e1003854.

117. Lee, C.R., Margolis, D.J. (2016). Pupil dynamics reflect behavioral choice and learning in a Go/NoGo tactile decision-making task in mice. Front Behav Neurosci, 10: 200.

118. Mathis, A., Mamidanna, P., Cury, K. M., Abe, T., Murthy, V. N., Mathis, M. W., Bethge, M. (2018). DeepLabCut: markerless pose estimation of user-defined body parts with deep learning. Nat Neurosci, 21: 1281–1289.

119. Nath, T., Mathis, A., Chen, A. C., Patel, A., Bethge, M., Mathis, M. W. (2019). Using DeepLabCut for 3D markerless pose estimation across species and behaviors. Nat Protocols, 14: 2152–2176.

120. Yamada, K., Toda, K. (2022). Pupillary dynamics of mice performing a Pavlovian delay conditioning task reflect reward-predictive signals. Front Syst Neurosci, 16.

121. Li, Y., Mathis, A., Grewe, B.F., Osterhout, J.A., Ahanonu, B., Schnitzer, M.J., Murthy, V.N., Dulac, C. (2017). Neuronal representation of social information in the medial amygdala of awake behaving mice. Cell, 171: 1176–1190.e17.

122. Kingsbury, L., Huang, S., Wang, J., Gu, K., Golshani, P., Wu, Y.E., Hong, W. (2019). Correlated neural activity and encoding of behavior across brains of socially interacting animals. Cell, 178(2): 429–446.e16.

123. Huda, R., Sipe, G.O., Breton-Provencher, V., Cruz, K.G., Pho, G.N., Adam, E., Gunter, L.M., Sullins, A., Wickersham, I.R., Sur, M. (2020). Distinct prefrontal top-down circuits differentially modulate sensorimotor behavior. Nat Commun, 11: 6007.

124. Ngyuen, C., Mondoloni, S., Le Borgne, T., Centeno, I., Come, M., Jehl, J., Solie, C., Reynolds, L.M., Durand-de Cuttoli, R., Tolu, S., et al. (2021). Nicotine inhibits the VTA-to-amygdala dopamine pathway to promote anxiety. Neuron, 109(16): 2604–2615.e9.

125. Mo, C., McKinnon, C., Sherman, S.M. (2023). A transthalamic pathway crucial for perception. BioRxiv, 10.1101/2023.03.30.533323.

126. Dallel, R., Raboisson, P., Auroy, P., Woda, A. (1988). The rostral part of the trigeminal sensory complex is involved in orofacial nociception. Brain Res, 448: 7–19.

127. Gauriau, C., Bernard, J.F. (2004). Posterior triangular thalamic neurons convey nociceptive messages to the secondary somatosensory and insular cortices in the rat. J Neurosci, 24: 752–761.

128. Hooks, B.M., Mao, T., Gutnisky, D.A., Yamawaki, N., Svoboda, K., Shepherd, G.M.G. (2013). Organization of cortical and thalamic input to pyramidal neurons in mouse motor cortex. J Neurosci, 33(2): 748–760.

129. Luo, P., Li, A., Zheng, Y., Han, Y., Tian, J., Xu, Z., Gong, H., Li, X. (2019). Whole brain mapping of long-range direct input to glutamatergic and GABAergic neurons in motor cortex. Front Neuroanat, 13(44).

130. Casas-Torremocha, D., Rubio-Teves, M., Hoerder-Suabedissen, A., Hayashi, S., Presna, L., Molnar, Z., Porrero, C., Clasca, F. (2022). A combinatorial input landscape in the “higher-order relay” posterior thalamic nucleus. J Neurosci, 42(41): 7757–7781.

131. Wilson, C.J. (1993). The generation of natural firing patterns in neostriatal neurons. Prog Brain Res, 99: 277–297.

132. Stern, E.A., Kincaid, A.E., Wilson, C.J. (1997). Spontaneous subthreshold membrane potential fluctuations and action potential variability of rat corticostriatal and striatal neurons in vivo. J Neurophysiol, 77: 1697–1715.

133. Reig, R., Silberberg, G. (2014). Multisensory integration in the mouse striatum. Neuron, 83: 1200–1212.

134. Mowery, T.M., Harrold, J.B., Alloway, K.D. (2011). Repeated whisker stimulation evokes invariant neuronal responses in the dorsolateral striatum of anesthetized rats: a potential correlate of sensorimotor habits. J Neurophysiol, 105: 2225–2238.

135. Cui, G., Jun, S., Jin, X., Pham, M.D., Vogel, S.S., Lovinger, D.M., Costa, R.M. (2013). Concurrent activation of striatal direct and indirect pathways during action inhibition. Nature, 494: 238–242.

136. Tecuapetla, F., Matias, S., Dugue, G.P., Mainen, Z.F., Costa, R.M. (2014). Balanced activity in basal ganglia projection pathways is critical for contraversive movements. Nat Commun, 5: 4315.

137. Klaus, A., Martins, G.J., Paixao, V.B., Zhou, P., Paninski, L., Costa, R.M. (2017). The spatiotemporal organization of the striatum encodes action space. Neuron, 95(5): 1171–1180.e7.

138. Day, M., Belal, M., Surmeier, W.C., Melendez, A., Wokosin, D., Tkatch, T., Clarke, V.R.J., Surmeier, D.J. (2024). GABAergic regulation of striatal spiny projection neurons depends upon their activity state. PLoS Biol, 22(1): e3002483.

139. Mastro, K.J., Bouchard, R.S., Holt, H.A., Gittis, A.H. (2014). Transgenic mouse lines subdivide external segment of globus pallidus (GPe) neurons and reveal distinct GPe output pathways. J Neurosci, 34: 2087–2099.

140. Baker, M., Kang, S., Hong, S., Song, M., Yang, M.A., Peyton, L., Essa, H., Lee, S.W., Choi, D. (2023). External globus pallidus input to the dorsal striatum regulates habitual seeking behavior in male mice. Nat Commun, 14(4085).

141. Reynolds, J.N.J., Avvisati, R., Dodson, P.D., Fisher, S.D., Oswald, M.J., Wickens, J.R., Zhang, Y. (2022). Coincidence of cholinergic pauses, dopaminergic activation, and depolarization of spiny projection neurons drives synaptic plasticity in the striatum. Nat Commun, 13: 1296.

142. Matityahu, L., Gilin, N., Sarpong, G.A., Atamna, Y., Tirohsi, L., Tritsch, N.X., Wickens, J.R., Goldberg, J.A. (2023). Acetylcholine waves and dopamine release in the striatum. Nat Commun, 14(6852).

143. Bokor, H., Frere, S.G., Eyre, M.D., Slezia, A., Ulbert, I., Luthi, A., Acsady, L. (2005). Selective GABAergic control of higher-order thalamic relays. Neuron, 45(6): 929–940.

144. Miller-Hansen, A.J., Sherman, S.M. (2022). Conserved patterns of functional organization between cortex and thalamus in mice. Proc Natl Acad Sci USA, 119(21): e2201481119.

145. Sachidhanandam, S., Sreenivasan, V., Kyriakatos, A., Kremer, Y., Petersen, C.C.H. (2013). Membrane potential correlations of sensory perception in mouse barrel cortex. Nat Neurosci, 16: 1671–1677.

146. Hemelt, M.E., Keller, A. (2008). Superior colliculus control of vibrissa movements. J Neurophysiol, 100: 1245–1254.

147. Schneider, G.E. (1969). Two visual systems. Science, 163: 895–902.

148. Ahmadlou, M., Zweifel, L.S., Heimel, J.A. (2018). Functional modulation of primary visual cortex by the superior colliculus in the mouse. Nat Commun, 9: 3895.

149. Klug, J.R., Engelhardt, M.D., Cadman, C.N., Li, H., Smith, J.B., Ayala, S., Williams, E.W., Hoffman, H., Jin, X. (2018). Differential inputs to striatal cholinergic and parvalbumin interneurons imply functional distinctions. eLife, 7: e35657.

150. Poppi, L.A., Ho-Nguyen, K.T., Shi, A., Daut, C.T., Tischfield, M.A. (2021). Recurrent implication of striatal cholinergic interneurons in a range of neurodevelopmental, neurodegenerative, and neuropsychiatric disorders. Cells, 10(907).

151. Dube, L., Smith, A.D., Bolam, J.P. (1988). Identification of synaptic terminals of thalamic or cortical origin in contact with distinct medium-size spiny neurons in the rat neostriatum. J Comp Neurol, 267: 455–471.

152. Wright, A.K., Norrie, L., Ingham, C.A., Hutton, E.A.M., Arbuthnott, G.W. (1999). Double anterograde tracing of outputs from adjacent “barrel columns” of rat somatosensory cortex. Neostriatal projection patterns and terminal ultrastructure. Neuroscience, 88: 119–133.

153. Raju, D.V., Ahern, T.H., Shah, D.J., Wright, T.M., Standaert, D.G., Hall, R.A., Smith, Y. (2008). Differential synaptic plasticity of the corticostriatal and thalamostriatal systems in an MPTP-treated monkey model of parkinsonism. Eur J Neurosci, 27(7): 1647–1658.

154. Hausser, M. (2001). Synaptic function: dendritic democracy. Curr Biol, 11(1): R10–R12.

155. Day, M., Wokosin, D., Plotkin, J.L., Tian, X., Surmeier, D.J. (2008). Differential excitability and modulation of striatal medium spiny neuron dendrites. J Neurosci, 28(45): 11603–11614.

156. Arai, R., Jacobowitz, D.M., Deura, S. (1994). Distribution of calretinin, calbindin D28k, and parvalbumin in the rat thalamus. Brain Res Bull, 35: 595–614.

157. Debanne, D. (2004). Information processing in the axon. Nat Rev Neurosci, 5: 304–316.

158. Citri, A., Malenka, R.C. (2008). Synaptic plasticity: multiple forms, functions, and mechanisms. Neuropsychopharmacol, 33: 18–41.

159. Kreitzer, A.C., Malenka, R.C. (2008). Striatal plasticity and basal ganglia circuit function. Neuron, 60(4): 543–554.

160. Lee, J.H., Durand, R., Gradinaru, V., Zhang, F., Goshen, I., Kim, D.S., Fenno, L.E., Ramakrishnan, C., Deisseroth, K. (2010). Global and local fMRI signals driven by neurons defined optogenetically by type and wiring. Nature, 465: 788–792.

161. Ranjan, R., Van Geit, W., Ruben, M., Rossert, C., Riquelme, J.L., Damart, T., Aurelien, J., Tuncel, I (2023). eFEL [computer software]. Zenodo, 10.5281/zenodo.593869.

162. Broussard, G.J., Liang, Y., Fridman, M., Unger, E.K., Meng, G., Xiao, X., Ji, N., Petreanu, L., Tian, L. (2018). In vivo measurement of afferent activity with axon-specific calcium imaging. Nat Neurosci, 21(1272).

163. Choung, A.S., Miri, M.L., Busskamp, V., Matthews, G.A.C., Acker, L.C., Sorensen, A.T., Young, A., Klapoetke, N., Henninger, M.A., Kodandaramaiah, S.B., et al. (2014). Noninvasive optical inhibition with a red-shifted microbial rhodopsin. Nat Neurosci, 17(8): 1123–1129.

164. Huber, D., Gutnisky, D.A., Peron, S., O’Connor, D.H., Wiegert, J.S., Tian, L., Oertner, T.G., Looger, L.L., Svoboda, K. (2012). Multiple dynamic representations in the motor cortex during sensorimotor learning. Nature, 484: 473–481.

165. Margolis, D.J., Lutcke, H., Schulz, K., Haiss, F., Weber, B., Kugler, S., Hasan, M.T., Helmchen, F. (2012). Reorganization of cortical population activity imaged throughout long-term sensory deprivation. Nat Neurosci, 15: 1539–1546.

166. Chen, J.L., Pfaffli, O.A., Voigt, F.F., Margolis, D.J., Helmchen, F. (2013). Online correction of licking-induced brain motion during two-photon imaging with a tunable lens. J Physiol, 591: 4869–4698.

167. Chen, J.L., Margolis, D.J., Stankov, A., Sumanovski, L.T., Schneider, B.L., Helmchen, F. (2015). Pathway-specific reorganization of projection neurons in somatosensory cortex during learning. Nat Neurosci, 18(8): 1101–1108.

168. Urai, A.E., Aguillon-Rodriguez, V., Laranjeira, I.C., Cazettes, F., The International Brain Laboratory, Mainen, Z.F., Churchland, A.K. (2021). Citric acid water as an alternative to water restriction for high-yield mouse behavior. eNeuro, 8(1): ENEURO.0230-20.2020.

169. Guo, Z.V., Hires, S.A., Li, N., O’Connor, D.H., Komiyama, T., Ophir, E., Huber, D., Bonardi, C., Morandell, K., Gutnisky, D., Peron, S., Xu, N., Cox, J., Svoboda, K. (2014). Procedures for behavioral experiments in head-fixed mice. PLoS One, 9: e88678.

170. Sanders, J.I., Kepecs, A. (2014). A low-cost programmable pulse generator for physiology and behavior. Frontiers in Neuroengineering, 7: 43.

171. McNicol, D. (1970). A primer of signal detection theory (Allen and Unwin).

172. Van der Walt, S., Schönberger, J. L., Nunez-Iglesias, J., Boulogne, F., Warner, J. D., Yager, N., Gouillart, E., Yu, T. (2014). Scikit-image: image processing in Python. PeerJ, 2: e453.

173. Legaria, A.A., Matikainen-Ankney, B.A., Yang, B., Ahanonu, B., Licholai, J.A., Parker, J.G., Kravitz, A.V. (2022). Fiber photometry in striatum reflects primarily nonsomatic changes in calcium. Nat Neurosci, 25: 1124–1128.

174. Oram T.B., Tenzer A., Saraf-Sinik I., Yizhar O., Ahissar E. (2024). Co-coding of head and whisker movements by both VPM and POm thalamic neurons. Nat Commun, 15(1):5883.

175. Petty G.H., Bruno R.M. (2024). Attentional modulation of secondary somatosensory and visual thalamus of mice. bioRxiv [Preprint] 2024.03.22.586242. doi: 10.1101/2024.03.22.586242. https://elifesciences.org/reviewed-preprints/97188

176. Smith Y., Galvan A., Ellender T.J., Doig N., Villalba R.M., Huerta-Ocampo I., Wichmann T., Bolam J.P. (2014). The thalamostriatal system in normal and diseased states. Front Syst Neurosci. 8:5.

177. Rodriguez-Moreno J, Porrero C, Rollenhagen A, Rubio-Teves M, Casas-Torremocha D, Alonso-Nanclares L, Yakoubi R, Santuy A, Merchan-Pérez A, DeFelipe J, Lübke JHR, Clasca F. (2020). Area-Specific Synapse Structure in Branched Posterior Nucleus Axons Reveals a New Level of Complexity in Thalamocortical Networks. J Neurosci. 40(13):2663–2679.

178. Koós T, Tepper JM. (1999). Inhibitory control of neostriatal projection neurons by GABAergic interneurons. Nat Neurosci. 2(5):467–72.

179. Fino E, Vandecasteele M, Perez S, Saudou F, Venance L. (2018). Region-specific and state-dependent action of striatal GABAergic interneurons. Nat Commun. 9(1):3339.

## Supplemental References

S1. Alloway, K.D., Smith, J.B., Mowery, T.M., Watson, G.D.R. (2017). Sensory processing in the dorsolateral striatum: the contribution of thalamostriatal pathways. Front Syst Neurosci, 11: 53.

S2. Alloway, K.D., Smith, J.B., Watson, G.D.R. (2014). Thalamostriatal projections from the medial posterior and parafascicular nuclei have distinct topographic and physiologic properties. J Neurophysiol, 111: 36-50.

S3. Mathis, A., Mamidanna, P., Cury, K. M., Abe, T., Murthy, V. N., Mathis, M. W., Bethge, M. (2018). DeepLabCut: markerless pose estimation of user-defined body parts with deep learning. Nat Neurosci, 21: 1281–1289.

S4. Nath, T., Mathis, A., Chen, A. C., Patel, A., Bethge, M., Mathis, M. W. (2019). Using DeepLabCut for 3D markerless pose estimation across species and behaviors. Nat Protocols, 14: 2152-2176.

S5. Van der Walt, S., Schönberger, J. L., Nunez-Iglesias, J., Boulogne, F., Warner, J. D., Yager, N., Gouillart, E., Yu, T. (2014). Scikit-image: image processing in Python. PeerJ, 2: e453.

S6. Lee, C.R., Margolis, D.J. (2016). Pupil dynamics reflect behavioral choice and learning in a Go/NoGo tactile decision-making task in mice. Front Behav Neurosci, 10: 200.

S7. Yamada, K., Toda, K. (2022). Pupillary dynamics of mice performing a Pavlovian delay conditioning task reflect reward-predictive signals. Front Syst Neurosci, 16.

